# Classifier architecture and data preprocessing jointly shape accelerometer-based behavioural inference

**DOI:** 10.64898/2026.02.16.706143

**Authors:** Loïc Brun, Jonas Rothrock, Erica van de Waal, Ebi Antony George

## Abstract

1. Although the use of accelerometer-based behavioural classification to quantify animal activity budgets is gaining widespread traction, the interactions between key preprocessing decisions and modern classification algorithms remain poorly understood. Moreover, classification pipelines are commonly assessed using global performance metrics, despite increasing evidence that such metrics poorly reflect behaviour-specific patterns and ecological reliability.
2. Using a free-ranging primate (*Chlorocebus pygerythrus*) as a case study, we benchmarked how temporal segmentation (burst length), collar orientation correction, and model architecture jointly shape behavioural inference. We compared nine supervised algorithms spanning classical machine learning, feature-based deep learning including a tabular foundation model (TabPFN), and state of the art time-series architectures (HydraMultiROCKET). Beyond conventional metrics, performance was further evaluated using ecological validation against independent focal observations to assess model stability and biological plausibility.
3. Model architecture exerted the strongest influence on classification outcomes. Modern deep-learning approaches substantially outperformed classical models, doubling recall for rare behaviours (e.g., grooming, self-scratching) without compromising precision. In contrast, burst length and collar orientation correction had little effect on global metrics but produced substantial, behaviour-specific trade-offs. Shorter bursts improved the detection of rare events by increasing training instances, while orientation correction suppressed dataset-specific artifacts at the cost of degrading common behaviours. Crucially, models with similar global and behaviour-level validation metrics produced divergent predictions when applied outside the annotated context.
4. Our findings reveal that global metrics are insufficient for optimizing behavioural inference in complex wild systems. We demonstrate that modern deep-learning architectures, such as the ROCKET family, provide a robust, accessible baseline that handles class imbalance more effectively than traditional methods. We propose that reliable inference requires behaviour-aware evaluation frameworks that integrate ecological validation, and advocate for ensemble or hierarchical strategies to leverage the complementary strengths of different preprocessing and modelling configurations.

## Introduction

Quantifying animal activity budgets is critical for understanding adaptive behavioural strategies and predicting species’ responses to environmental change (Berger-Tal et al., 2011; Dunbar et al., 2009). Advances in biologging technologies have transformed this task by enabling the collection of high-resolution, time-series data on animal movement and behaviour (Cooke et al., 2004). Among these tools, tri-axial accelerometers are particularly powerful, allowing fine-scale behavioural inferences without direct observation (Nathan et al., 2012). However, accelerometers generate large, complex datasets whose analysis remains methodologically challenging (Hoffman et al., 2024). Increasingly, these challenges are addressed using machine learning (ML), which is well suited to detect nonlinear, multidimensional structure in behavioural data (Valletta et al., 2017). Across taxa, the reliability of accelerometer-based behavioural inference is shaped by three methodological stages: (i) logger deployment and cross-context variability, (ii) data preprocessing, and (iii) behavioural inference. While deployment-related variation (e.g. tag placement, attachment orientation and sampling regime) is often system specific and difficult to control (Brown et al., 2013; Garde et al., 2022), preprocessing and algorithm choice are more amenable to optimization and standardization. We therefore dissect how these latter two stages jointly shape behavioural inference using accelerometer data from vervet monkeys (*Chlorocebus pygerythrus*) as a case study.

A core preprocessing choice is temporal segmentation whereby acceleration data is partitioned into fixed-length windows for behavioural classification. This choice entails a fundamental trade-off as short windows may fail to capture the temporal structure of a behaviour, whereas long windows may increase the likelihood of aggregating multiple behaviours within a single segment. Reported optimal window durations vary widely across taxa and behavioural scales, ranging from sub-second windows for rapid behaviours in birds (Yu et al., 2023) to windows exceeding one minute in mammals (Tatler et al., 2018). Burst length therefore directly shapes class representation, label purity, and ultimately classification performance.

Another persistent challenge is sensor orientation. Collar-mounted loggers can rotate or shift over time, altering the mapping between sensor axes and body orientation and reducing the transferability of behavioural classifiers across time or individuals (Garde et al., 2022; Kamminga et al., 2018). Orientation-invariant metrics, such as Euclidean norm of acceleration, can partially mitigate this issue but do so by removing axis-specific information that may be pertinent for behavioural discrimination. These metrics have been shown to underperform orientation-dependent features even under deliberate tag rotation (Versluijs et al., 2023). The use of additional sensors like gyroscopes or magnetometers incur higher power costs and have yielded inconsistent or marginal benefits in wild deployments compared to accelerometers alone (Chakravarty et al.,2019; Conners et al., 2021). An alternative approach addresses sensor rotation by estimating sensor tilt (roll and pitch) from the static acceleration component of the accelerometer signal, enabling partial correction of collar orientation (Rautiainen et al., 2022). However, because gravity provides no information about rotation around the vertical axis, yaw remains unresolved without an external directional reference. In species capable of actively manipulating collars, such as primates, this unresolved orientation variability may further degrade inference. In such scenarios, using the accelerometer data itself to virtually align the sensor with the animal’s body represents a promising, yet untested, avenue for recovering fine-scale behavioural kinematics.

Other preprocessing decisions, including behavioural class definition and feature engineering may influence classification outcomes. Although these factors can substantially affect performance (e.g. Christensen et al., 2023; Fehlmann et al., 2017) a systematic evaluation is beyond the scope of this study; we therefore focus on burst length and sensor orientation, and critically, their interaction.

The final stage of accelerometer-based inference concerns algorithm choice. Behavioural classification has traditionally relied on classical machine-learning approaches trained on engineered features, such as Random Forests (RF), or gradient-boosted trees (XGBoost), which perform well across taxa and datasets (Mauny et al., 2025; Nathan et al., 2012; Rast et al., 2020; Yu et al., 2021). However, these models operate on summary representations of each data segment and do not explicitly model the temporal ordering of acceleration within segments. This may limit their ability to capture fine-scale or sequential behaviours and reduce transferability across individuals or contexts (Bidder et al., 2014; Rast et al., 2020; Wilson et al., 2018).

In contrast, deep-learning architectures designed for time-series data can learn hierarchical and temporal representations directly from raw acceleration signals, bypassing feature engineering and showing promising performance in comparative evaluations of ecological accelerometry data (Hoffman et al., 2024; Otsuka et al., 2024). However, their application often requires massive datasets and extensive hyperparameter tuning, which has limited their uptake in wildlife studies where labelled data are often sparse and imbalanced. Recent architectures are beginning to overcome these constraints: computationally efficient time-series classifiers such as the ROCKET family (Dempster et al., 2023) perform competitively with minimal tuning, and data-efficient tabular foundation models (Hollmann et al., 2025) are explicitly designed to generalise well under limited data regimes. Together, these advances lower the technical and data-related barriers that have historically constrained the adoption of deep learning in ecological accelerometry by combining computational efficiency, reduced tuning requirements, and accessible implementations.

Despite these advances, it remains unclear how preprocessing choices interact with model architecture to shape behavioural inference in complex systems. We address this question using a free-ranging primate, the vervet monkey, a system in which frequent transitions between terrestrial and arboreal locomotion, and complex social interactions constitute a stringent test case. Using tri-axial acceleration data, we classified eight behaviours (resting, sleeping, eating, walking, running, grooming receiver, grooming actor, and self-scratching) and conducted four complementary experiments: (i) testing how burst length influences classification performance and stability, (ii) evaluating the effects of orientation correction using an accelerometer-only rotation framework, (iii) comparing nine supervised classification algorithms spanning classical ML, feature-based DL, and time-series DL architectures, and (iv) validating accelerometer-derived activity budgets against independent focal observations. By explicitly linking preprocessing decisions, algorithmic architecture, and ecological validation, our study provides practical guidance for improving the reliability and interpretability of accelerometer-based behavioural inference in complex systems.

## Methods

### Study site and collar deployment

This study was conducted on seven habituated groups (20-80 individuals per group) of wild vervet monkeys at the INKAWU Vervet Project (IVP), located in the Mawana Game Reserve, KwaZulu-Natal, South Africa (28°00.327′S, 031°12.348′E). In June 2022, 37 adult individuals were fitted with UHF 1C Light collars (e-obs GmbH, Germany). The collar deployment was part of a larger study on male dispersal. Logger settings were optimized for deployment period of 1.5 years, and tri-axial acceleration was recorded at 10 Hz in bursts of 13.8 s every 90 s until December 2023 (Fig. 1A). All the raw accelerometer data is stored and accessible on Movebank (Study ID: 2111088601; Kays et al., 2022).

**Figure 1:**
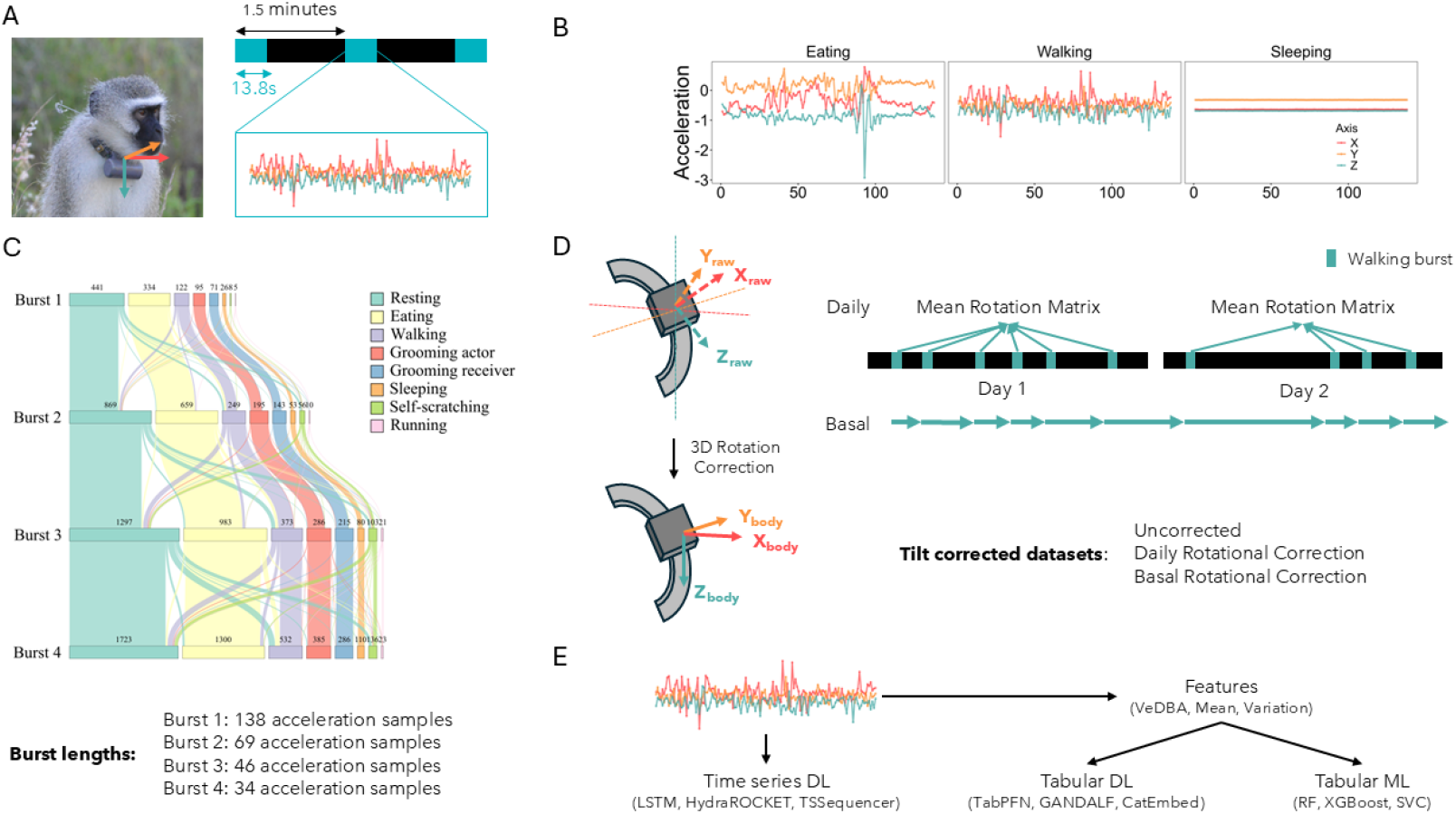
Data structure, preprocessing steps, and classification framework. **(A)** Collar-mounted tri-axial accelerometer and burst-sampling scheme. Acceleration was sampled at 10 Hz in bursts of 13.8 s (indicated in turquoise) every 90 s. **(B)** Example of tri-axial acceleration signals for three representative behaviours (eating, walking, and sleeping); colours indicate the three accelerometer axes (X, Y, Z). **(C)** Behavioural composition of annotated data across the four burst-length conditions. Original 13.8 s bursts (Burst 1) were subdivided into windows of 6.9 s (Burst 2), 4.6 s (Burst 3), and 3.4 s (Burst 4), altering the relative representation of behaviours. **(D)** Schematic of the collar orientation correction. Raw sensor axes were rotated into a body-centred reference frame using walking bouts to estimate a forward axis from the dynamic acceleration component, with gravity defining the vertical axis. Two correction schemes were applied: a daily mean rotation and a basal correction based on a rolling window mean. **(E)** Classification pipelines. Raw acceleration time series were used directly for time-series deep-learning models, whereas features were extracted for tabular deep-learning and machine-learning models.

### Ethical Statement

Captures for collar deployment were conducted by a certified veterinarian, following protocols approved by the University of KwaZulu-Natal (UKZN) Animal Ethics Committee and by the European Research Council, in accordance with Horizon Europe regulations. EvdW is affiliated to UKZN and collaborates with Prof. Collen Downs who has been granted animal ethics permission to capture, fit loggers and release vervet monkeys with the assistance of a veterinarian (application AREC/00005320/2023(00020574), Project title: Ecosystem health and biodiversity: The ecology, physiology, behaviour and conservation of selected southern African vertebrates). Only individuals (6 females, 31 males) for whom the collar mass represented < 3% of total body mass were selected for deployment. At the end of the study, another capture session took place to remove the collars.

### Video recording and annotation

To generate a labelled training dataset, we recorded 158 videos (43 h total) from 21 collared individuals between August 2022 and July 2023. Videos were annotated in BORIS (Friard & Gamba, 2016) using an ethogram with 39 behaviours/behavioural contexts (Table S1) by LB and JR, who calibrated their scoring through joint training sessions to ensure consistency. While most behaviours were mutually exclusive, up to three concurrent codes were allowed to capture composite activities (e.g., self-scratching while chewing and foraging). Rare events like aggression were recorded opportunistically, resulting in an unbalanced video dataset across individuals (mean = 124 min, SD = 91, range: 17–355 min) and behavioural classes (mean = 70 min, SD=122, range: 0.26-494 min).

### Temporal alignment of acceleration samples and video observations

Video and acceleration data were synchronized in Firetail (Berger et al., 2022), resulting in a final training dataset of 1,102 bursts (4.24 h). The severely reduced training dataset is due to the non-continuous burst-sampling protocol (one burst of 13.8 s every 90 s containing 138 acceleration samples) and the removal of some videos with uncertain temporal alignment (n = 5). Each acceleration sample was then assigned a behavioural label based on the video annotation. “Pure” bursts containing a single behaviour were assigned directly, whereas multi-behaviour bursts were labelled using a majority rule, with the assigned class corresponding to the behaviour represented by most acceleration samples. Given the limited number of behaviours that could be reliably distinguished from collar-mounted tri-axial accelerometers, we collapsed the original 39 annotated behaviours into 8 main categories (Fig. 1B, Table S1). This was done based on kinematic similarity (e.g. sitting resting and sitting vigilant), shared biological states (e.g. climbing up and climbing down) and biological relevance (e.g. grooming actor vs. grooming receiver). The resulting distribution of acceleration bursts across behavioural categories and individuals was highly uneven, reflecting both behavioural prevalence and sampling constraints (Fig. S1).

#### Experiment 1: Effect of burst length

Most machine learning classifiers require a single output, here one behaviour, per burst. However, a 13.8 s burst (comprising 138 samples), often captures multiple behaviours, introducing uncertainty in label assignment. Because shorter bursts are expected to reduce the proportion of multi-behaviour segments and improve the representation of short-lived behaviours (e.g. self-scratching; Fig. S2), we hypothesized that reducing burst duration would improve classification accuracy. To test this, we systematically subdivided the original 13.8 s burst windows (burst 1) into progressively shorter durations of 6.9 s, 4.6 s, and 3.4 s (burst 2, burst 3, burst 4 respectively; Fig. 1C). We compared model and behaviour-level classification performance across these burst lengths.

#### Experiment 2: Effect of collar orientation correction

Collar orientation can shift over time and might differ among individuals. Under ecologically realistic conditions with imbalanced datasets, this variation in sensor tilt can bias learning toward orientation-specific signals rather than behaviour-related kinematics and consequently, reduce model transferability across contexts and individuals. For instance, in our dataset, sleeping exhibited strong individual bias in collar orientation (Fig. S3). To mitigate this issue, we implemented a three-dimensional body-frame alignment that accounts for rotation around all three axes (roll, pitch, and yaw). Gravity-based alignment methods use the static acceleration component to correct sensor orientation relative to gravity, thereby resolving pitch and roll but leaving rotation around the vertical (yaw) axis unresolved (e.g. Rautiainen et al., 2022). Therefore, to estimate the animal’s full anatomical frame, we used stereotyped walking bouts to infer the forward body axis. Walking bursts were identified from the full dataset using smoothed vectorial dynamic body acceleration (VeDBA) with a threshold of 0.17–0.30 (Fig. S4), leveraging the relatively consistent kinematics of forward locomotion across individuals. For each burst, the vertical body axis was defined using the mean static acceleration vector, corresponding to gravity. To resolve rotation around this axis (yaw), dynamic acceleration was projected onto the horizontal plane, and the forward axis was defined as the first principal component of the horizontal dynamic acceleration, assuming that the direction of maximum variance corresponds to the direction of travel during locomotion. The lateral axis was then obtained as the cross-product of the vertical and forward vectors, yielding a right-handed, orthonormal body-centred reference frame that was subsequently applied to the full time series.

Orientation correction was applied at two scales (Fig. 1D): daily correction (averaged across all walking bouts in 24 h), and (ii) basal correction (rolling average of recent bouts, with a window size of 20, 40, 60, and 80 bouts for bursts 1 to 4 respectively). We compared model and behaviour-level classification performance across three datasets: uncorrected, with daily corrections and with basal corrections.

#### Experiment 3: Comparison of classification algorithms

To assess how algorithmic framework and input representation influence classification performance, we compared nine supervised models (Fig. 1E; model descriptions and full names in Table S2) spanning three methodological groups: (i) feature-based ML algorithms (RF, XGBoost, SVM), (ii) feature-based DL algorithms (CEM, GANDALF, TabPFN), and (iii) time-series DL algorithms (LSTM, TSSequencer, HydraMultiROCKET). Feature-based models utilized 60 variables derived from established time-and frequency-domain approaches (e.g. Fehlmann et al., 2017; Nathan et al., 2012), including descriptive statistics, dynamic acceleration metrics (e.g., VeDBA), spectral descriptors, and PCA-based features (Table S3). Time-series models were trained directly on the raw acceleration signal. We employed a stratified 75:25 train:validation split, repeated five times to ensure stability, yielding 540 models (9 algorithms × 3 datasets × 4 burst lengths × 5 repeats). Hyperparameters were kept at default settings to facilitate standardized comparison, with early stopping applied to deep learning models.

#### Experiment 4: Validation against traditional focal sampling

We compared model-derived activity budgets against independent focal observations (Altmann, 1974) based on the ethogram in Table S1 and collected daily as part of the long-term monitoring at IVP. To minimize seasonal effects, we selected the three months with the highest sampling effort across collared individuals, yielding 400 focal follows from 32 individuals (mean duration = 18 min; mean 12.5 follows/individual). Focal data were compared with predictions from the two best-performing models (from Experiment 3) applied to seven randomly selected days per individual within the same period. Both datasets were mapped to the same eight behavioural categories. The best performing models were trained on the burst 1 and burst 4 dataset, yielding 511,052 and 2,044,214 predicted acceleration samples respectively (burst 1: mean ± SD = 15970 ± 5083 samples per individual, range = 2495 – 20146; burst 4: mean ± SD = 63882 ± 20331 samples per individual, range = 9980 – 80584). For the activity budget comparison, model predictions were restricted to daytime hours (06:00–18:00) to match focal sampling effort. Additionally, we assessed consistency in circadian activity patterns by visually comparing hourly proportions of active (walking, running, grooming actor, self-scratching, eating) and inactive (grooming receiver, resting and sleeping) behaviours across 24-hours.

### Statistical Analysis

To simplify analysis, behaviours were categorized by their prevalence in the ground truth dataset: common (> 15% of the dataset; Resting, Eating), uncommon (< 15% and > 5%; Walking, Grooming actor, Grooming receiver) and rare (< 5%; Sleeping, Self-scratching, Running). We evaluated global model performance using Receiver Operating Characteristic Area Under the Curve (ROC AUC) and accuracy, and behaviour-specific performance using precision and recall.

In experiments 1–3, we used Generalized Linear Models (GLMs) for global performance comparisons and Generalized Linear Mixed-effects Models (GLMMs) for behavioural-level analyses. All models utilized a beta error distribution, with response variables transformed to the open interval (0, 1) when necessary (Smithson & Verkuilen, 2006). GLMMs included a random intercept for seed. The single fixed predictor in the GLMs corresponded to the experimental manipulation: burst length (experiment 1, four levels), correction type (experiment 2, three levels), or model architecture (experiment 3, nine levels). For behavioural-level analyses using GLMMs, we first tested for interactions between the experiment-specific predictor and behavioural category using likelihood ratio tests. Non-significant interactions (experiment 1 and 2) were followed by analysis of main effects and comparisons of the estimated marginal means between levels of the predictors independently, while significant interactions (experiment 3) were followed up using post-hoc comparisons of estimated marginal means conditional on behaviour. Baseline models for each experiment were selected to isolate the variable of interest. Experiment 1 used uncorrected RF models across burst lengths to test the effect of burst duration. Experiment 2 fixed burst length to burst 1 and used RF models to test the effect of collar orientation correction. Experiment 3 used the uncorrected burst 1 dataset to compare algorithmic architectures. In experiment 1, rare behaviours were excluded from behavioural-level analyses due to insufficient sample size in burst 1 while in experiments 2 and 3, the analyses were conducted using burst 4, which provided sufficient representation of rare behaviours. In experiment 3, we further examined model performance across all eight behaviours classes instead of the three behavioural categories. Since global comparisons yielded similar results for ROC AUC and accuracy, we report ROC AUC in the main text and accuracy in the Supplementary Material.

In experiment 4, we compared activity budgets using GLMMs with a beta error distribution. The response variable was the proportion of time allocated to each behaviour, modeled with predictors for data source (Focal vs. Model 1 vs. Model 2) and behavioural class. Individual identity was included as a random intercept to account for repeated measures. We first tested for an interaction between data source and behavioural class. Given a significant interaction between predictors, post-hoc comparisons assessed differences between data sources within each behavioural class using estimated marginal means.

All the statistical analysis were done in R version 4.3.2 (R Core Team, 2023). We used the glmmTMB (Brooks et al., 2017) package to fit the statistical models, DHARMa (Hartig, 2024) and performance (Lüdecke et al., 2021) packages to validate model assumptions, emmeans (Lenth & Piaskowski, 2025) and modelbased (Makowski et al., 2025) packages to estimate marginal means and perform posthoc comparisons and the tidyverse (Wickham et al., 2019) package for data handling and plotting.

Machine learning and deep learning models were trained using python versions 3.10 and 3.12 (Van Rossum & Drake, 2009). We used the sklearn (Pedregosa et al., 2011) package for RF and SVM models, xgboost (Chen & Guestrin, 2016) package for XGBoost, PyTorch Tabular (Joseph, 2021) package for the Category Embedding Model and GANDALF, the tabpfn (Hollmann et al., 2025) package for TabPFN and the tsai (Oguiza, 2023) package for LSTM, TSSequencer and HydraMultiROCKET models. In addition, we used numpy (Harris et al., 2020) and pandas (McKinney, 2010) packages for data handling.

## Results

### Experiment 1: Effect of burst length

Burst length had no significant effect on overall model performance as measured by ROC AUC (Fig. 2A; χ^2^ = 1.5, p = 0.68; Table S4). All pairwise comparisons among burst lengths were non-significant (p > 0.05 for all; Table S5;). At the behaviour level, rare behaviours were excluded from the analysis due to insufficient sample size in burst 1. Comparing common and uncommon behaviours, we found no significant interaction between burst length and behaviour type for either precision (χ^2^ = 4.16, p = 0.25) or recall (χ^2^ = 0.58, p = 0.90). However, behaviour type itself significantly predicted performance (precision: χ^2^ = 12.42, p < 0.001; recall: χ^2^ = 80.31, p < 0.001) while burst length did not (p > 0.05 for main effects of burst length). Uncommon behaviours had higher precision, but lower recall than common behaviours (Fig. 2C; Table S6; precision = 0.73 – 0.78 vs. 0.67 – 0.72; recall=0.51 – 0.54 vs. 0.79 – 0.81; p < 0.001 for all comparisons; Table S7).

**Figure 2:**
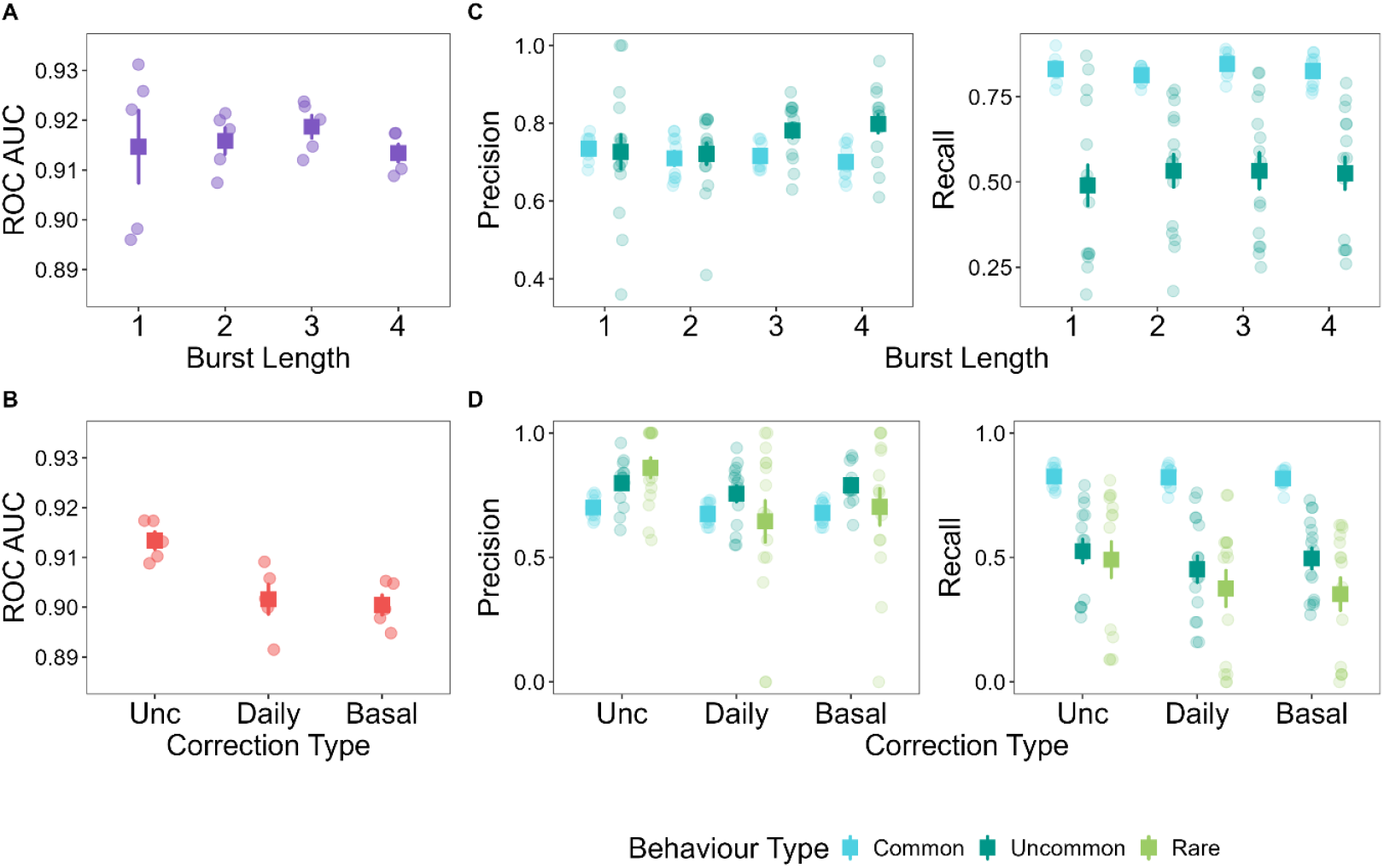
Effects of burst length and collar orientation correction on Random Forest (RF) performance. **(A)** Overall classification performance measured as ROC AUC across burst lengths for models trained on uncorrected data. **(B)** ROC AUC across orientation correction types (uncorrected, daily, basal) for models trained on burst 1 data. **(C)** Behaviour-level precision and recall for common and uncommon behaviours across burst lengths. **(D)** Behaviour-level precision and recall for common, uncommon, and rare behaviours across orientation correction types (burst 4 data). Circles represent results from five independent training runs using different random seeds and larger squares indicate the mean across runs, with error bars showing the 95% confidence intervals in all panels.

### Experiment 2: Effect of collar orientation correction

The orientation correction had a significant effect on the ROC AUC of RF models trained on the burst 1 dataset (χ^2^ = 8.77, p = 0.01) but did so in a direction opposite to our hypothesis. Both the daily and basal rotational correction had similarly lower ROC AUC compared to the uncorrected dataset (Fig. 2B; z = -2.60 for comparison of both correction types with the uncorrected dataset, p = 0.009; z = -0.001 for daily vs basal correction comparison, p = 0.99; Table S5).

At the behaviour level, correction type showed no significant interaction with behaviour type for either precision (χ^2^ = 8.30, p = 0.08) or recall (χ^2^ = 3.99, p = 0.41). However, correction type significantly affected precision but not recall (precision: χ^2^ = 7.79, p = 0.02; recall: χ^2^ = 5.00, p = 0.08), whereas behaviour type affected recall and not precision (precision: χ^2^ = 4.95, p = 0.08; recall: χ^2^ = 67.01, p < 0.001). The daily correction had significantly lower precision than the uncorrected dataset (Fig. 2D, Tables S6 and S8; daily correction: 0.57 – 0.69; uncorrected: 0.71 – 0.80; z = -2.75, p = 0.006) while the basal rotational correction had similar precision to both the daily rotational correction dataset and the uncorrected dataset (basal correction: 0.62 – 0.73; p > 0.05 for comparisons with uncorrected and daily correction). Across behavioural types, the pattern in recall was the same as observed in experiment 1. Recall was highest for common behaviours, intermediate for uncommon behaviours and was the lowest for rare ones (Tables S6 and S7; common: 0.73 – 0.80; uncommon: 0.45 – 0.55; rare: 0.31 – 0.40; p < 0.001 for all pairs of comparisons).

### Experiment 3: Comparison of classification algorithms

Model algorithm significantly affected the ROC AUC of models trained on the burst 1 uncorrected dataset (Fig. 3; χ^2^= 1345.43, p < 0.001). HydraMultiROCKET emerged as the top-performing model, achieving a mean ROC AUC of 0.95 (CI: 0.94-0.96; Table S4). Crucially, pairwise comparisons confirmed this performance was statistically superior to every other model tested (Table S5; all p < 0.05). The next-best model, TabPFN, attained a significantly lower ROC AUC of 0.93 (CI: 0.92-0.94; Table S4). Traditional ML models RF and XGBoost showed comparable performance to each other (ROC AUC ∼ 0.91; Table S4) but were outperformed by the top two models. At the other extreme, the sequence-specific DL model, LSTM, performed poorly (ROC AUC: 0.57; Table S4) and was significantly outperformed by all other methods (all pairwise p < 0.001; Table S5).

**Figure 3:**
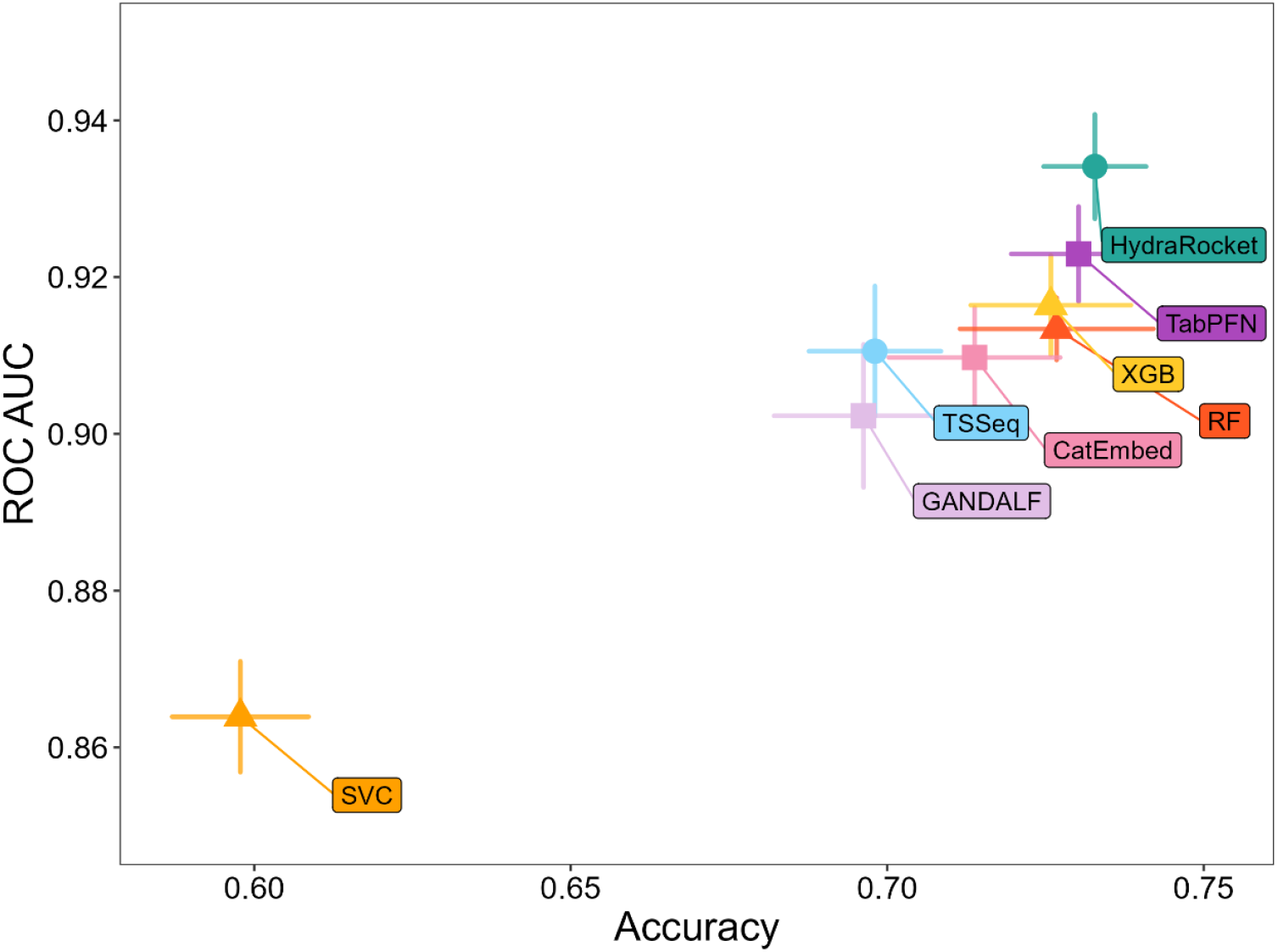
Comparison of classification performance across models. The accuracy and ROC AUC of models trained on burst 1 uncorrected data is plotted. Points represent the mean performance across five independent training runs using different random seeds, with horizontal and vertical error bars indicating the 95% confidence intervals for accuracy and ROC AUC, respectively. The LSTM model (mean accuracy = 0.40, mean ROC AUC = 0.61) is omitted from the plot to preserve scale and highlight differences.

Using the burst 4 dataset, the analysis at the behavioural level revealed a significant interaction effect between the model algorithm and behaviour rarity for both precision (χ^2^= 64.24, p < 0.001) and recall (χ^2^ = 64.05, p < 0.001). We focused on the distinct performance profiles of the three top-performing, representative models of each type: RF, TabPFN, and HydraMultiROCKET. The RF model exhibited a clear precision-recall trade-off driven by class imbalance. Its precision scaled with rarity (Table S6; common: 0.65, CI:0.51–0.79; uncommon: 0.73, CI: 0.63–0.83; rare: 0.88, CI: 0.83–0.94), with rare behaviours showing significantly higher precision than common and uncommon behaviours (Table S7; rare vs common: p = 0.02; rare vs uncommon: p = 0.03). This came at the cost of a reduction in recall for all non-common behaviours (Tables S6 and S7; common: 0.76, CI: 0.66–0.87 vs. uncommon: 0.52, CI: 0.41–0.63 and rare: 0.47, CI: 0.36– 0.58; p < 0.001 for both uncommon and rare vs. common).

In contrast, the deep-learning models, TabPFN and HydraMultiROCKET, showed a more stable profile. Unlike RF, their precision remained statistically consistent across all rarity levels (Table S6; TabPFN = common: 0.65, CI: 0.52–0.79; uncommon: 0.71, CI: 0.61–0.82;rare: 0.71, CI: 0.61–0.82; HydraMultiROCKET = common: 0.67, CI: 0.53–0.80; uncommon:0.68, CI: 0.57–0.78; rare: 0.79, CI: 0.70–0.87; all pairwise comparisons p>0.05). While both DL models experienced a drop in recall for non-common behaviours similar to RF, they still maintained higher sensitivity in these difficult classes (Table S6; TabPFN = common: 0.75, CI: 0.65–0.86 vs. uncommon: 0.54, CI: 0.44–0.65 and rare: 0.57, CI: 0.46–0.68; HydraMultiROCKET = common: 0.75, CI: 0.64–0.86; uncommon: 0.57, CI: 0.46–0.67 and rare: 0.60, CI: 0.49–0.70; p < 0.05 for both uncommon and rare vs. common). Crucially, HydraMultiROCKET achieved the most balanced profile as it combined the highest overall ROC AUC, strong statistical stability in overall precision and the highest absolute performance metrics for rare behaviours (Precision: 0.79; Recall: 0.60; Table S6), distinguishing it as the most reliable model for capturing underrepresented behaviours.

Finally, to identify the single best-performing model for each behaviour, we compared the top-performing configurations across all three selected models, burst lengths, and correction types for precision and recall. This analysis confirmed that deep learning models (TabPFN and HydraMultiROCKET) outperformed the best machine learning model as they were the top model in 13 of the 16 categories (Fig. 4; 2 metrics of precision and recall across 8 behaviours). HydraMultiROCKET and TabPFN were equally good, with 6 and 7 best-performing slots and similar precision and recall values in several behaviours. However, HydraMultiROCKET performed better with common behaviours, especially in the case of precision. On the other hand, TabPFN performed better for rare behaviours like Running and Sleeping. We also detected effects of burst length and orientation correction on behaviour-specific performance. Burst 1 and burst 4 were the best performing window lengths in 5 categories each, with the former predominantly performing the best in common behaviours like Resting and Eating and the latter performing better for rare behaviours like Sleeping and Running. The effect of the correction type was less pronounced with the clearest effect in Sleeping, where the basal correction with burst 4 was the best performing model by a large margin (Figs. 4 and S5). Accordingly, the two most dominant models, which performed the best in each of 3 categories, were HydraMultiROCKET trained on burst 1 data with the uncorrected dataset and TabPFN trained on burst 4 data with the basal correction.

**Figure 4:**
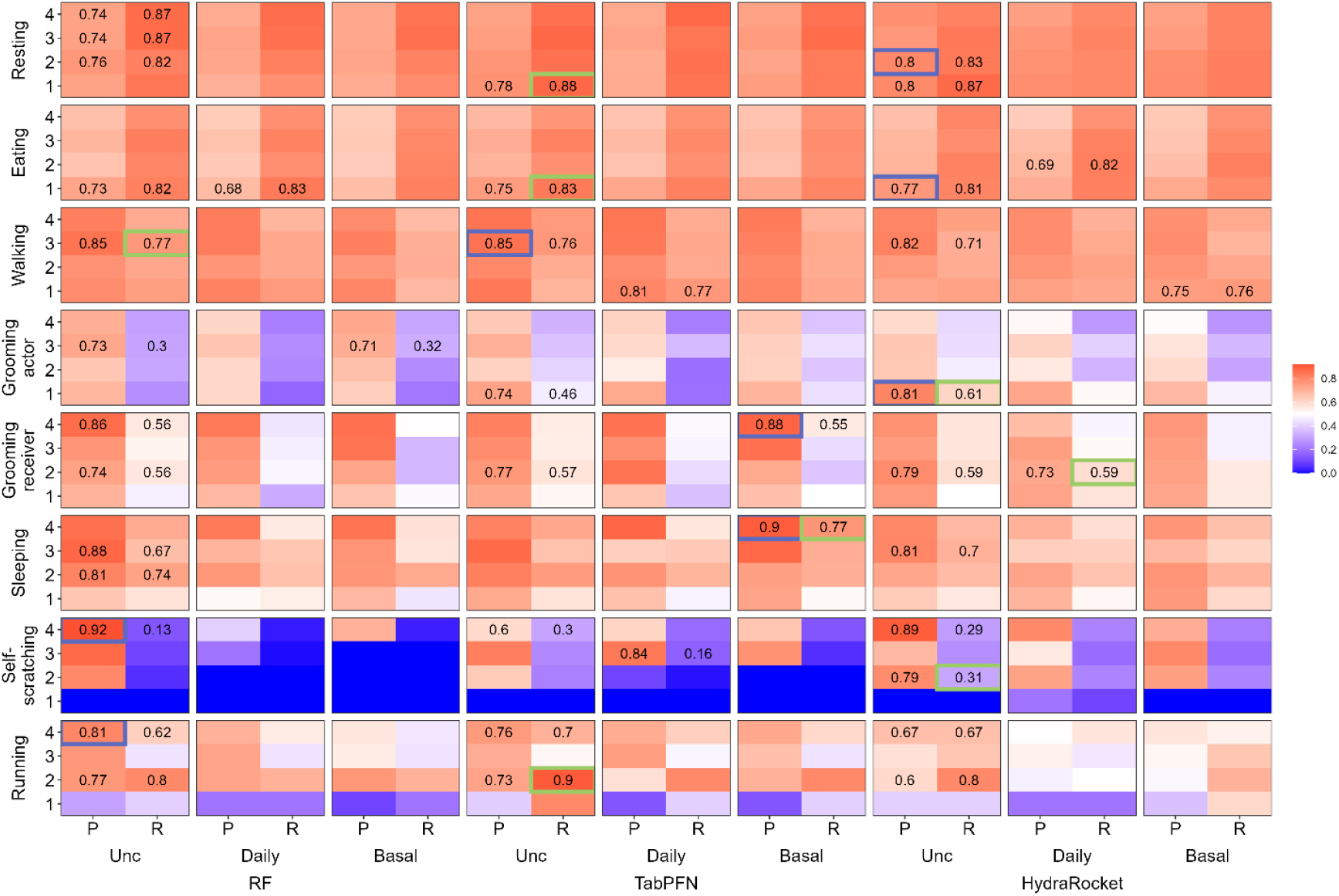
Comparison of behaviour-specific performance of the top models. The precision (P) and recall (R) for Random Forest, TabPFN, and HydraMultiROCKET across orientation correction types and burst lengths is plotted. The colour intensity of the tiles indicates performance (blue = low, red = high). For each behaviour and model algorithm, only the values of the specific models with the highest precision and recall are provided within tiles. When these metrics were achieved by different models, values of both models are reported. Across behaviour, the specific model with the highest precision and recall are further highlighted using a purple and green border around the tile respectively.

### Experiment 4: Comparison against traditional focal sampling

Overall, activity budgets derived from the predictions from the two best-performing models (HydraMultiROCKET and TabPFN) reflected the same general behavioural patterns as the focal data, while providing fine-grained estimates with reduced variance (Fig. 5A). The statistical comparison revealed a significant interaction effect between the measurement source (focal vs. model predictions) and behaviour type (χ^2^ = 400, p < 0.001). Interestingly, both models predicted a lower mean proportion of Eating and Self-scratching compared to focal sampling data while predicting a higher proportion of the six other behaviours. Comparing the two DL models, the proportion of behaviours predicted differed in the case of Resting, Grooming receiver and Self-scratching with the TabPFN predictions showing a higher proportion of Resting and Self-scratching and a lower proportion of Grooming receiver (Tables S9 and S10; Resting: 39.31% vs 32.86%, z= 4.18, p < 0.001; Self-scratching: 2.36% vs 1.15%, z = 4.50, p < 0.001; Grooming receiver: 7.49% vs 10.18%, z = -3.22, p = 0.001). The predicted proportion for all other five behaviours were not significantly different between the two models (Table S10; p > 0.05 for all 5 comparisons). Aggregation into active and inactive behaviours yielded similar circadian profiles across models (Fig. 5B), whereas behaviour-specific temporal patterns showed greater divergence, especially for Resting, Grooming receiver, and Sleeping (Fig. S6).

**Figure 5:**
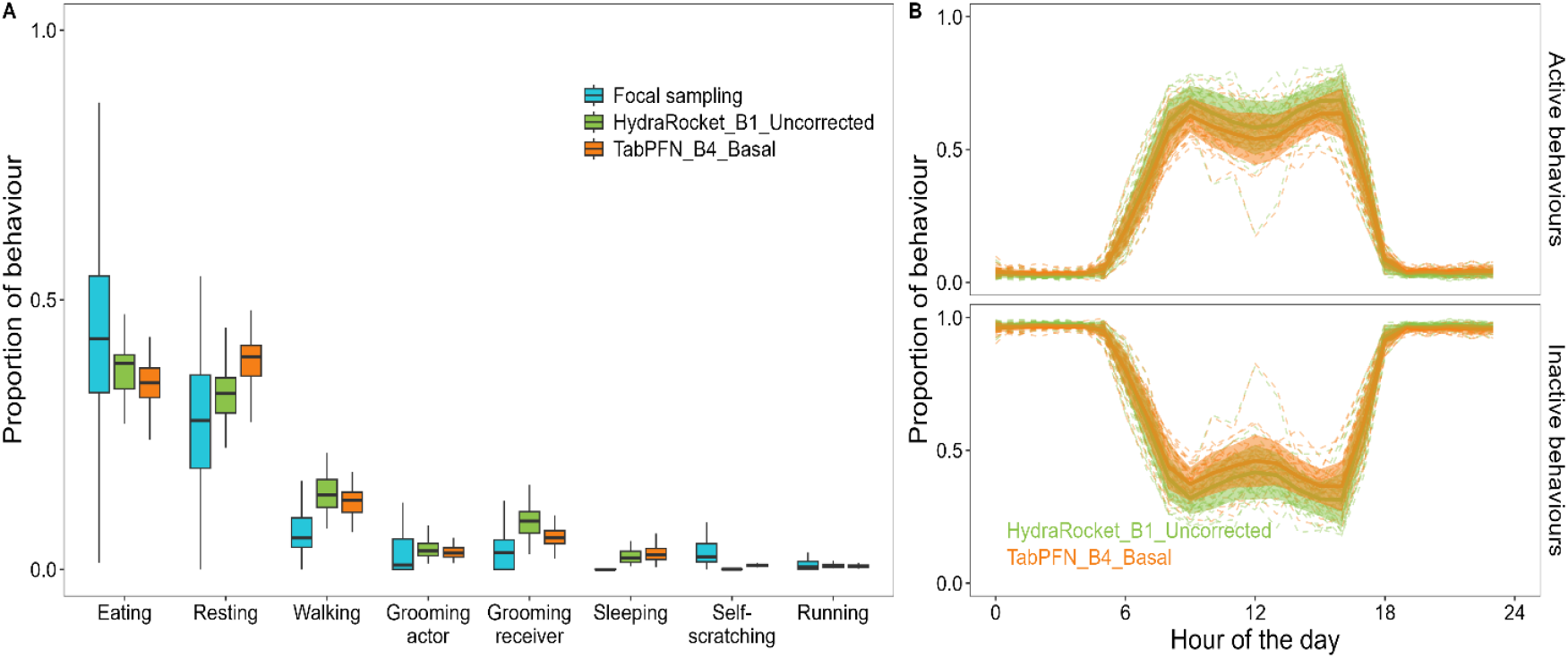
Ecological validation of classification performance. **(A)** Behavioural proportions from focal sampling and models predictions from the HydraMultiROCKET model trained on uncorrected burst 1 dataset and TabPFN model trained on burst 4 dataset with basal orientation correction. Box plots represent the inter-quartile range of the distribution of proportions across all individuals and months with the median provided by the horizontal black line. The error bars represent the range of the distribution excluding outliers. The boxplots are coloured based on the source of measurement of the proportions, with blue for focal sampling, green for the HydraMultiROCKET model and orange for the TabPFN model. **(B)** Circadian patterns of active and inactive behaviours predicted by the two models. Dotted lines represent data from each individual averaged over all the days of predictions for that individual, while the solid line and the shaded region represent the mean and standard deviation around the mean for the entire group. The lines are coloured based on the model as in A.

## Discussion

We examined how two key preprocessing choices, temporal segmentation (burst length) and collar orientation correction, interact with algorithmic architecture to shape accelerometer-based behavioural inference in a free-ranging primate. Algorithm choice emerged as the dominant driver of classification performance, with modern DL architectures consistently outperforming traditional ML models. In contrast, burst length and orientation correction had limited effects on overall metrics. Crucially, however, behaviour-specific analyses and ecological validation of model predictions revealed that this apparent lack of effect concealed significant differences for underrepresented behaviours. Together, our results demonstrate that preprocessing and modelling decisions should not be evaluated independently and that global performance metrics alone are insufficient for assessing ecological reliability.

### Algorithmic architecture outweighs preprocessing in complex behavioural systems

Modern DL approaches clearly outperformed classical ML models, but these gains were architecture-specific rather than universal to deep learning, in line with recent benchmark studies (Hoffman et al., 2024; Otsuka et al., 2024). Time-series convolutional methods from the ROCKET family (HydraMultiROCKET) and a pretrained tabular foundation model (TabPFN) consistently outperformed RF and XGBoost, whereas a standard recurrent DL architecture (LSTM) performed poorly under default settings. In our system, characterized by limited training data, strong class imbalance, and short discontinuous acceleration bursts, these two models succeeded through complementary mechanisms. HydraMultiROCKET bypasses explicit feature engineering and large parameter optimization, which constrain recurrent DL models, by relying on large banks of random convolutional kernels (Dempster et al., 2023). This makes it exceptionally efficient for classifying short time-series segments where data is insufficient to train complex recurrent weights (Reimers & Gurevych, 2017). In contrast, TabPFN operates on engineered features but leverages statistical priors learned from massive synthetic datasets, to achieve strong generalization on small, imbalanced tabular data (Hollmann et al., 2025). It effectively brings the foundation model paradigm which has revolutionized fields such as natural language processing and computer vision to tabular datasets like feature-based accelerometer classification (Bommasani et al., 2022; Brown et al., 2020). Crucially, both architectures are available in ready-made python libraries that enable easy usage and can be deployed with minimal tuning, offering a robust alternative to classical ML models without the engineering complexity typically associated with deep learning.

The main advantage of these models was not only higher overall performance, but a significantly more balanced error distribution across behavioural classes. While classical models like RF achieved high precision for dominant classes, they suffered from poor sensitivity to underrepresented behaviours. In contrast, HydraMultiROCKET and TabPFN models substantially improved recall for rare classes without compromising precision. For example, when comparing the best classical and deep-learning models, recall doubled for grooming actor (0.32 to 0.61) and self-scratching (0.13 to 0.31), with only minor decreases in precision for self-scratching (0.92 to 0.89) and running (0.81 to 0.76; Fig. 4). This demonstrates that modern DL architectures can recover rare, ecologically critical behaviours that are typically masked by global performance metrics in imbalanced datasets.

### Burst length impacts class imbalance rather than global performance

While burst length had no detectable effect on global performance metrics (ROC AUC, accuracy), it significantly influenced model stability (Fig. 2). Models trained on longer burst durations (burst 1) exhibited greater variability across random seeds, whereas shorter burst durations (burst 4) yielded more stable performance. This instability likely stems from the reduced sample size and increased probability of mixed-behaviour segments in longer windows (Fig. S2), which introduces label ambiguity and makes performance highly sensitive to specific data splits.

At the behavioural level, shorter bursts improved the detection of rare behaviours while longer bursts favoured common classes (Fig. 4). Importantly, this effect was independent of the intrinsic duration of a behaviour, as rare behaviours included both brief, high-activity events (e.g. running) and longer, low-activity states (e.g. sleeping), with high label purity irrespective of burst length (Fig. S2). This suggests that shorter bursts primarily benefit rare classes by increasing the number of training instances available to the model rather than by isolating short events. Conversely, for common behaviours where sample size is not a limiting factor, longer bursts allow the model to learn from longer temporal dependencies, resulting in higher performance. Consequently, optimal burst length represents a trade-off between maximizing sample size for rare classes and capturing temporal context for common ones.

### Orientation correction can remove biologically informative structure

In collar-mounted wildlife accelerometry and in the absence of additional sensors, variation in orientation is typically handled indirectly using orientation-invariant features (Bidder et al., 2014; Kamminga et al., 2018; Shepard et al., 2008), and less commonly through partial orientation correction based on the gravity component of static acceleration to recover roll and pitch (Christensen et al., 2023; Rautiainen et al., 2022). We tested a novel accelerometer-only approach estimating yaw from stereotyped walking bouts. Contrary to expectations, this global correction generally reduced classification performance. The failure likely stems from the violation of our core assumption that locomotion always occurs along a single linear axis while walking. When animals engage in circular movement or complex vertical displacement, or when valid walking bouts are absent between collar shifts, the inferred reference frame becomes unstable. Under these conditions, reprojection introduces additional variance rather than reducing noise. This aligns with previous work showing that orientation correction can substantially alter the structure of inertial sensor signals used for behaviour classification, with effects that are not uniformly beneficial across behaviours or analytical workflows (Chakravarty et al., 2019; Conners et al., 2021; Garde et al., 2022).

However, behaviour-specific analyses revealed a critical exception as performance substantially increased for sleeping but decreased for common static behaviours (e.g. resting) and for rare yet highly dynamic behaviours (e.g. running). This improvement does not reflect enhanced kinematic recovery, but rather the removal of a sampling artifact. Our sparse sleeping data were inadvertently associated with specific collar orientations. Orientation correction neutralized this spurious dependency, preventing the model from exploiting static tilt as a shortcut and forcing it to learn intrinsic behavioural features (Fig.S3). Our results are in line with recent work demonstrating that increasing orientation variability through random rotation or data augmentation can improve classification of rare behaviours by decreasing reliance on axis-specific signal structure (Otsuka et al., 2024). In this context, imperfect yaw estimation may act as a weak form of regularization for under-represented behaviours, while degrading performance for other classes by introducing additional variance through unstable orientation estimation or by removing biologically meaningful posture information. This mechanism is illustrated by increased dispersion of static acceleration values under basal correction and reduced dispersion under daily correction for resting (Fig. S7) Since global performance metrics cannot distinguish between the removal of biologically meaningful signal and the suppression of dataset-specific artefacts (Wilson et al., 2025), behaviour-specific evaluation is essential to determine whether apparent performance gains arise from spurious associations with dataset artefacts or from genuine behavioural learning.

### Ecological validation reveals limits of metric-based evaluation

Comparison of model-derived activity budgets with independent focal observations confirmed the ecological relevance of our behavioural inference pipeline. Daytime predictions closely matched focal data, with inter-individual variability in predicted behaviour exceeding differences between the two shortlisted models (Fig. 5). However, while HydraMultiROCKET and TabPFN shared similar validation metrics, they yielded distinct proportion estimates for specific behaviours (e.g., resting). This discrepancy highlights the difficulty in directly translating traditional performance metrics into biologically meaningful inferences in wild systems characterized by limited training data. In contrast, nighttime predictions exhibited substantially greater divergence between models (Fig. S6). Both models predicted an extensive proportion of Grooming receiver during the night, which is biologically implausible given that vervet monkeys are strictly diurnal. However, the magnitude of this error varied between models. Indeed, TabPFN, selected for its superior performance on sleeping, showed markedly less confusion between sleeping and grooming receiver than HydraMultiRocket. These departures from biological expectations are most plausibly explained by limitations in the training data, as sleeping was sparsely represented and annotated exclusively from daytime video recordings. Importantly, sleep posture differs between daytime and nighttime, with seated positions more common during daytime naps and recumbent postures predominating at night. The latter can kinematically resemble grooming receiver postures, increasing the risk of misclassification (based on field observations; LB). Consequently, models validated on daytime data failed when behavioural expression shifted across temporal contexts, further emphasizing the necessity of biological validation beyond standard performance metrics (Christensen et al., 2025).

Finally, both models consistently underestimated eating and self-scratching while slightly overestimating other behaviours compared to focal sampling. This reflects differences in temporal resolution and behavioural definitions rather than just classification bias. First,focal sampling exhibited high inter-individual variance due to limited sampling effort, whereas high-resolution accelerometer data minimized this variation. Second, despite efforts to harmonize annotations, the methods differ in context. Real-time focal sampling records broad ecological contexts (e.g., ‘foraging’ includes walking and pausing), whereas models classify discrete kinematic states (e.g., walking, resting, eating) based on post-hoc video annotation. Furthermore, brief signals like eating are often masked by dominant locomotor signals within a burst. Thus, behaviours defined by social or ecological context often lack unique kinematic signatures distinct from static postures, like the similarity between receiving grooming and resting, making them prone to misclassification without supplementary contextual data.

### Recommendation and next steps

Our results support a shift in how accelerometer-based inference is approached in behavioural ecology. First, researchers should actively consider modern deep learning approaches, particularly ROCKET-family time-series methods and tabular foundation models, as these require minimal tuning yet deliver robust performance even with limited, imbalanced data. However, these tools must be applied with a focus on ecological utility rather than global optimization. We found that, in line with previous studies (Garde et al., 2022; Otsuka et al., 2024), standard metrics often mask substantial trade-offs, where configurations with similar overall model performance differed in classifying rare, biologically critical behaviours. Consequently, model selection and preprocessing (e.g., burst length) should be treated as behaviour-dependent design decisions. This should be complemented with the reporting of behaviour-specific metrics to ensure that performance aligns with the study’s biological questions rather than arbitrary benchmarks.

Second, because different architectures and preprocessing settings excel for different behaviours, we recommend moving away from single-model pipelines toward ensemble or hierarchical classification strategies. Ensemble approaches have already proven effective in accelerometer-based livestock behaviour classification (Wang et al., 2018) and a similar logic can be readily adapted to wildlife accelerometry. Strategies such as hierarchical branching (e.g., classifying overall activity level followed by behavioural sub-classes) or voting between models are promising avenues for further exploration. Such approaches allow researchers to leverage the complementary strengths of different methodologies, for example improving sensitivity to rare events while retaining robustness for stable states. Ultimately, future advances will likely depend less on identifying a single best classifier and more on workflows that integrate predictions across models and rigorously validate them against biological expectations.

## Supporting information

This manuscript contains one additional supplementary PDF containing Figures S1 to S7 681 and Tables S1 to S10.

## Author contributions

L.B.: Conceptualization, Methodology, Software, Validation, Formal Analysis, Investigation, Data Curation, Writing - Original Draft, Writing - Review & Editing, Visualization; J.R.: Investigation, Data Curation; E.vd.W.: Conceptualization, Funding Acquisition, Resources, Supervision, Writing—Review & Editing; E.A.G.: Conceptualization, Methodology, Software, Validation, Formal Analysis, Investigation, Data Curation, Writing - Original Draft, Writing - Review & Editing, Visualization.

## Acknowledgements

We thank the van der Walt family for permission to conduct this study on their land. We are deeply grateful to the entire INKAWU Vervet Project (IVP) team for their assistance in the field and during logger deployment, and especially to the on-site manager, Siboniso Thela, for his help in locating and retrieving dispersing individuals. We also thank the field assistants, Veronica Khosana, Lou Coudurier and Zonke Mbutho, for their invaluable support during data collection. We are particularly grateful to Michel Halbwax, the veterinarian, for his outstanding commitment during the deployment and retrieval of the loggers, as well as to Mike Henshall and Maria Granell Ruiz for their extensive assistance during logger deployment.

## Data availability statement

All the code and data required to replicate the results of this manuscript are available through Zenodo (https://doi.org/10.5281/zenodo.18486872). This also includes detailed instructions on how to set up the python environments and run the training of different models discussed in this manuscript.

## Supporting Information

This manuscript contains one additional supplementary PDF containing Figures S1 to S7 and Tables S1 to S10.

