## Supplementary material for "Classifier architecture and data preprocessing jointly shape accelerometer-based behavioural inference": This manuscript contains one additional supplementary PDF containing Figures S1 to S7 681 and Tables S1 to S10.

We used ROC AUC and accuracy to compare the global performance of the models in experiments 1 – 3. Since we obtained similar results for ROC AUC and accuracy, we provide results for the ROC AUC in the main manuscript and for accuracy here.

#### Experiment 1: Effect of burst length

Burst length had no significant effect on overall model accuracy (χ² = 2.9, p = 0.41). All pairwise comparisons among burst lengths were non-significant (p > 0.05 for all).

#### Experiment 2: Effect of collar orientation correction

The orientation correction had a significant effect on the accuracy of RF models trained on the burst 1 dataset (χ^2^ = 8.23, p = 0.02). However, the effect was the opposite of our hypothesis. Both the daily and basal rotational correction had similarly lower accuracy compared to the uncorrected dataset (z = -2.5 for comparison of both correction types with the uncorrected dataset, p = 0.01; z = 0.04 for daily vs basal correction comparison, p = 0.97).

#### Experiment 3: Comparison of classification algorithms

Model algorithm significantly affected the accuracy of models trained on the burst 1 uncorrected dataset (χ^2^ = 515.17, p < 0.001). HydraMultiROCKET emerged as the top-performing model, achieving a mean accuracy of 0.77 (CI: 0.74-0.79). Crucially, pairwise comparisons confirmed this performance was statistically superior to every other model tested except TabPFN (all p < 0.05; Table S1). The next-best model, TabPFN, attained a slightly lower accuracy of 0.76 (CI: 0.74-0.79). Traditional ML models RF and XGB showed comparable performance to each other (accuracy ~ 0.73) but were outperformed by HydraMultiROCKET and to a lesser extent by TabPFN. At the other extreme, sequence-specific models like LSTM performed poorly with an accuracy of 0.40 (CI: 0.37-0.43) and was significantly outperformed by all other methods (all pairwise p < 0.001; Table S1).

### Supplementary Figures

##### Figure S1

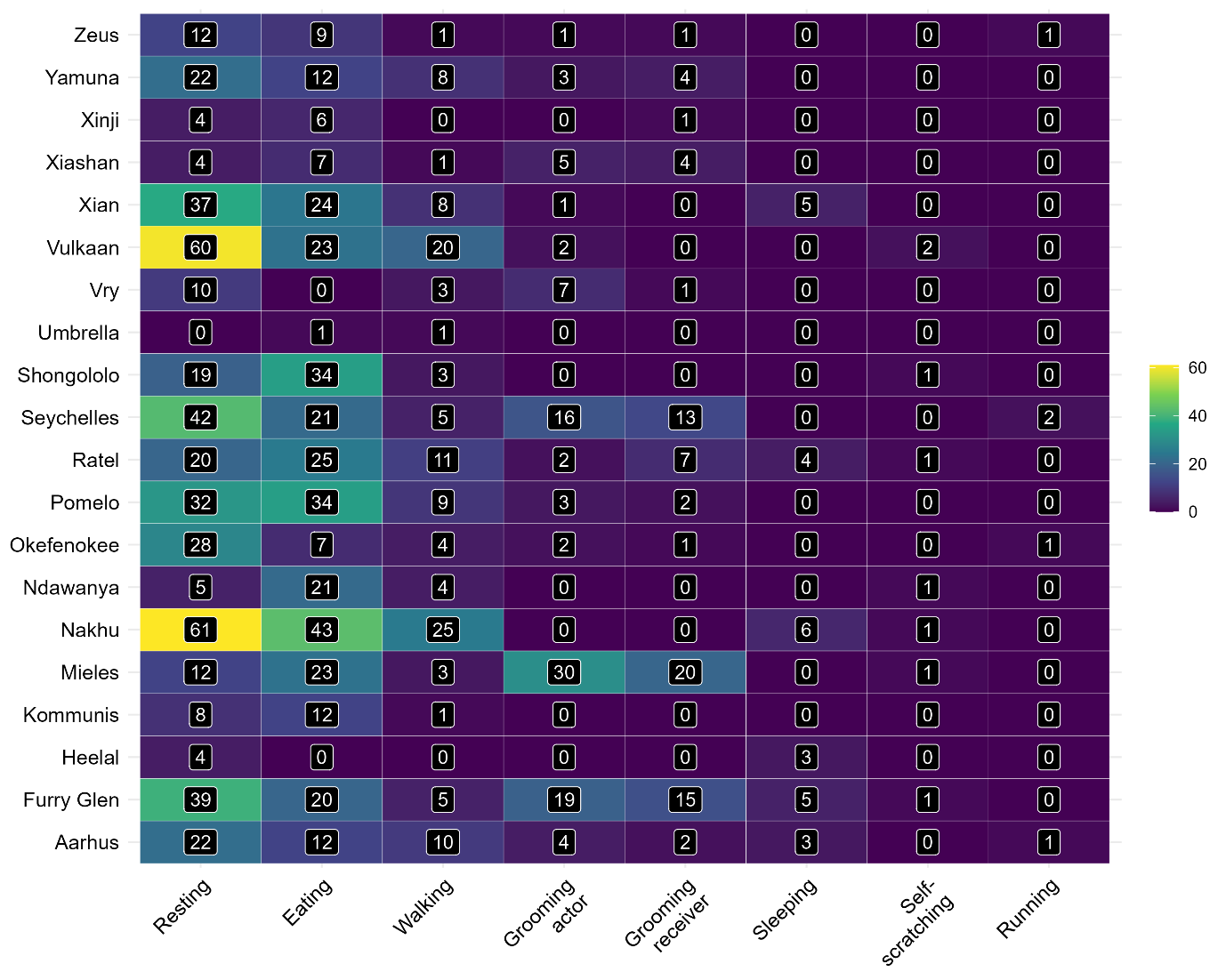

**Figure S1:** Number of acceleration bursts in the burst 1 dataset per behaviour and individual. The tiles are coloured by the number of samples and the numbers inset into each tile gives the actual value.

##### Figure S2

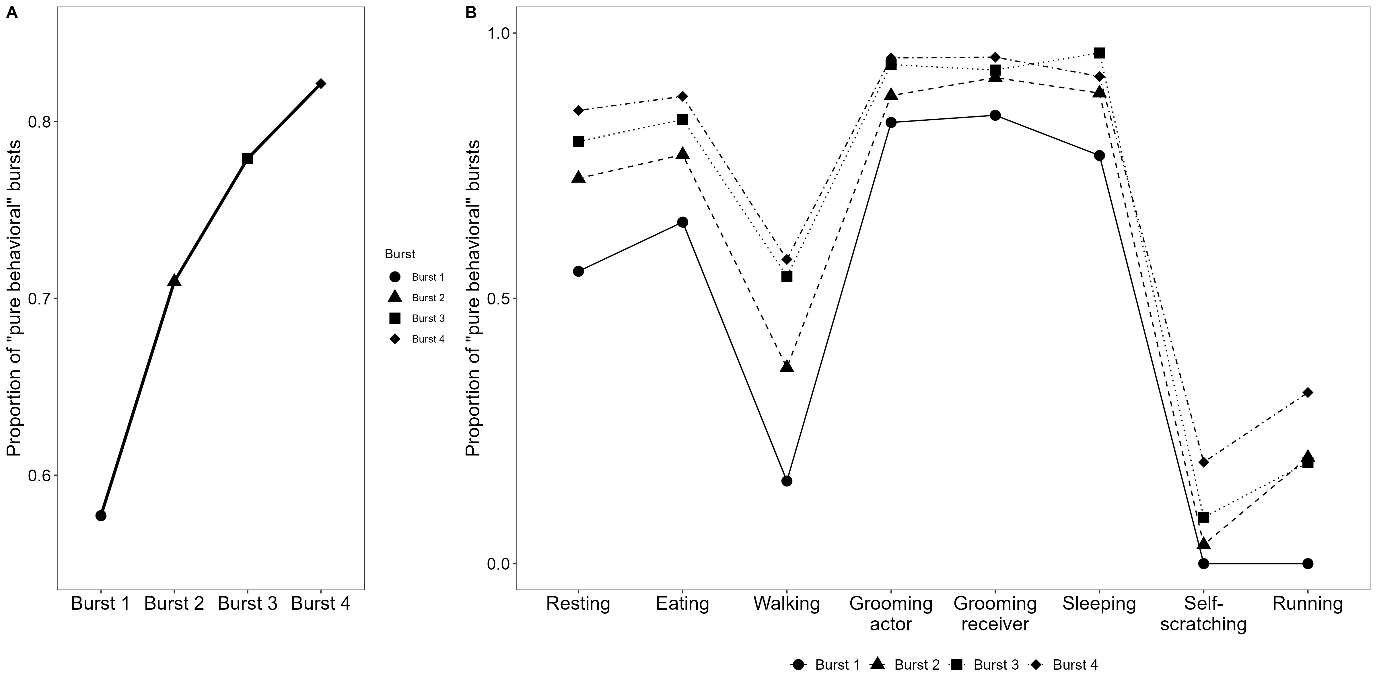

**Figure S2**: A) The proportion of “pure behavioural” bursts (i.e., bursts in which all acceleration samples were assigned to the same behaviour) with changing burst length. Different shapes denote different burst lengths. B) The proportion of “pure behavioural” bursts per behavioural class. Different shapes and line types denote different burst lengths.

##### Figure S3

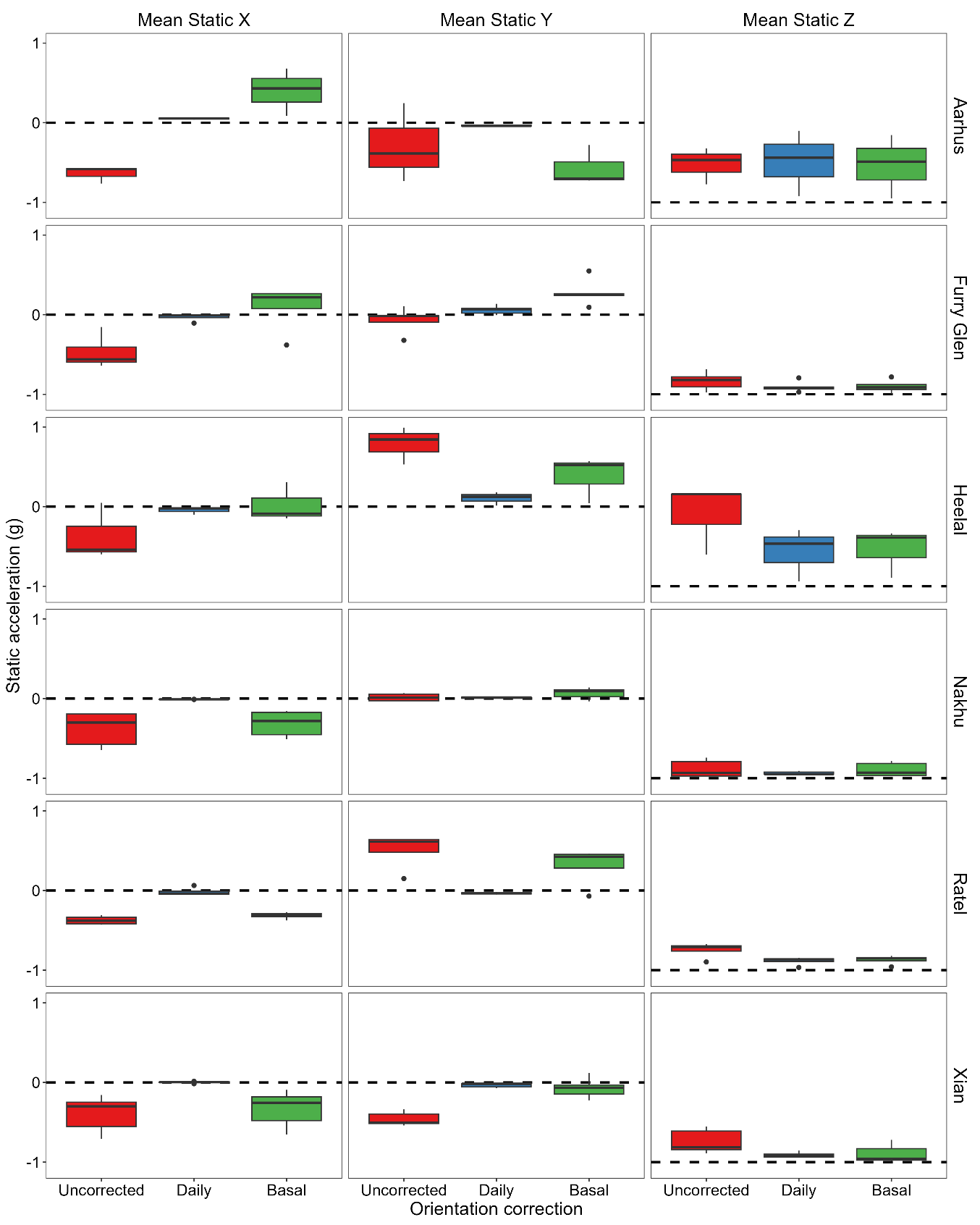

**Figure S3**: The mean static acceleration of X, Y and Z axes during bursts classified as sleeping from the raw uncorrected dataset and the daily and basal 3D orientation corrected datasets. Each facet represents data from one individual and one axis. Orientation correction defines a body-cantered reference frame based on walking bouts, with the Z-axis aligned to gravity and the horizontal axes defined relative to the estimated forward direction during locomotion. Under this reference frame, static acceleration during walking is expected to be close to 0 on the X and Y axes and −1 on the Z axis when alignment is stable. Deviations from these values during sleeping reflect differences between walking posture and sleeping posture, as well as residual variability in orientation estimation. Apparent lateral tilts observed in some individuals (e.g. Ratel versus Xian on the Y axis) likely reflect uneven representation of sleeping postures in the training data rather than intrinsic behavioural differences. Boxplots are coloured by the correction type.

##### Figure S4

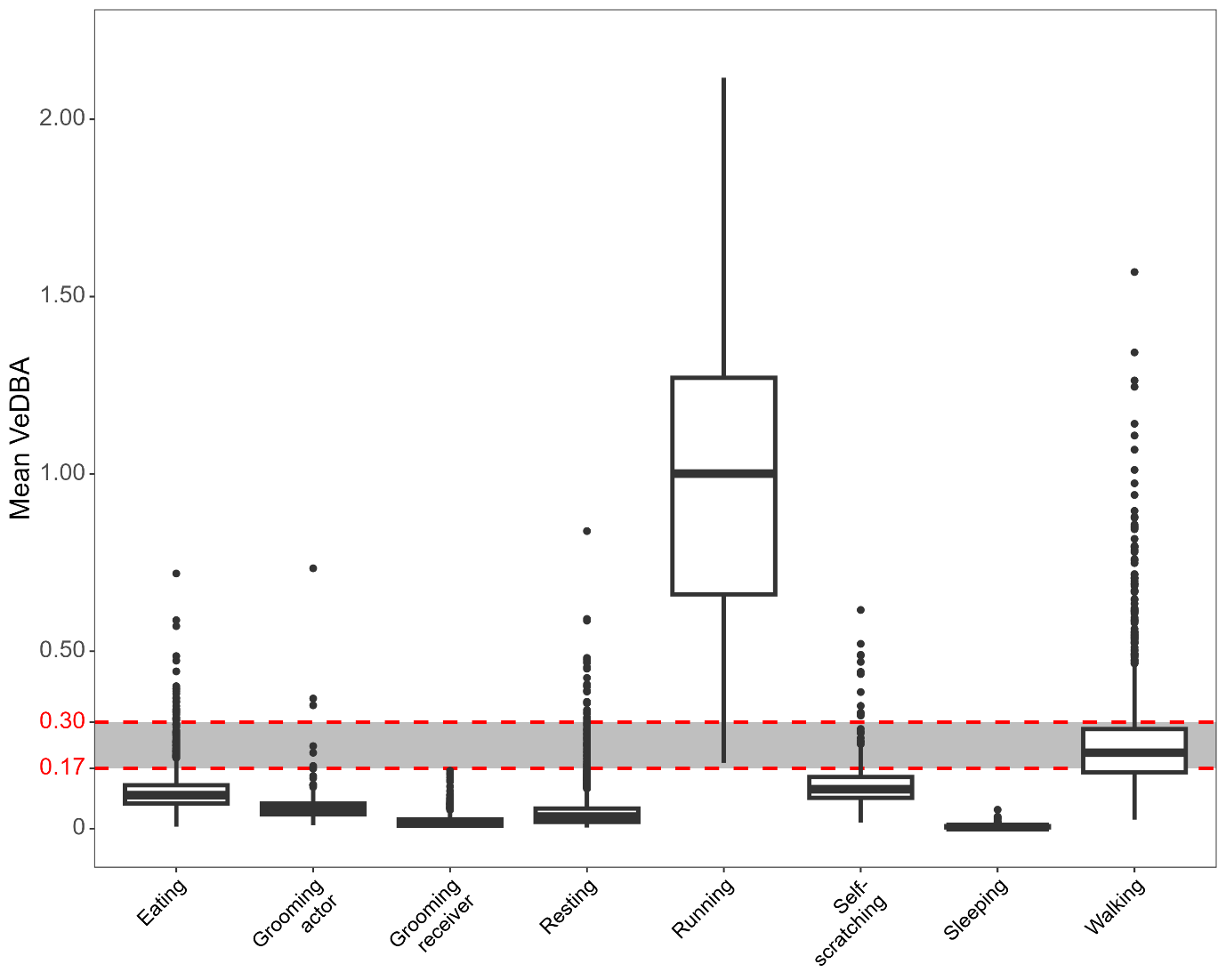

**Figure S4**: The mean VeDBA of acceleration bursts from each annotated behavioural class across all four burst length datasets. The threshold used to subset walking bursts for the orientation correction is highlighted as a grey rectangle. Box plots represent the inter-quartile range, horizontal lines inset within the boxplots represent the median, the whiskers represent the whole distribution and circles represent outliers.

##### Figure S5

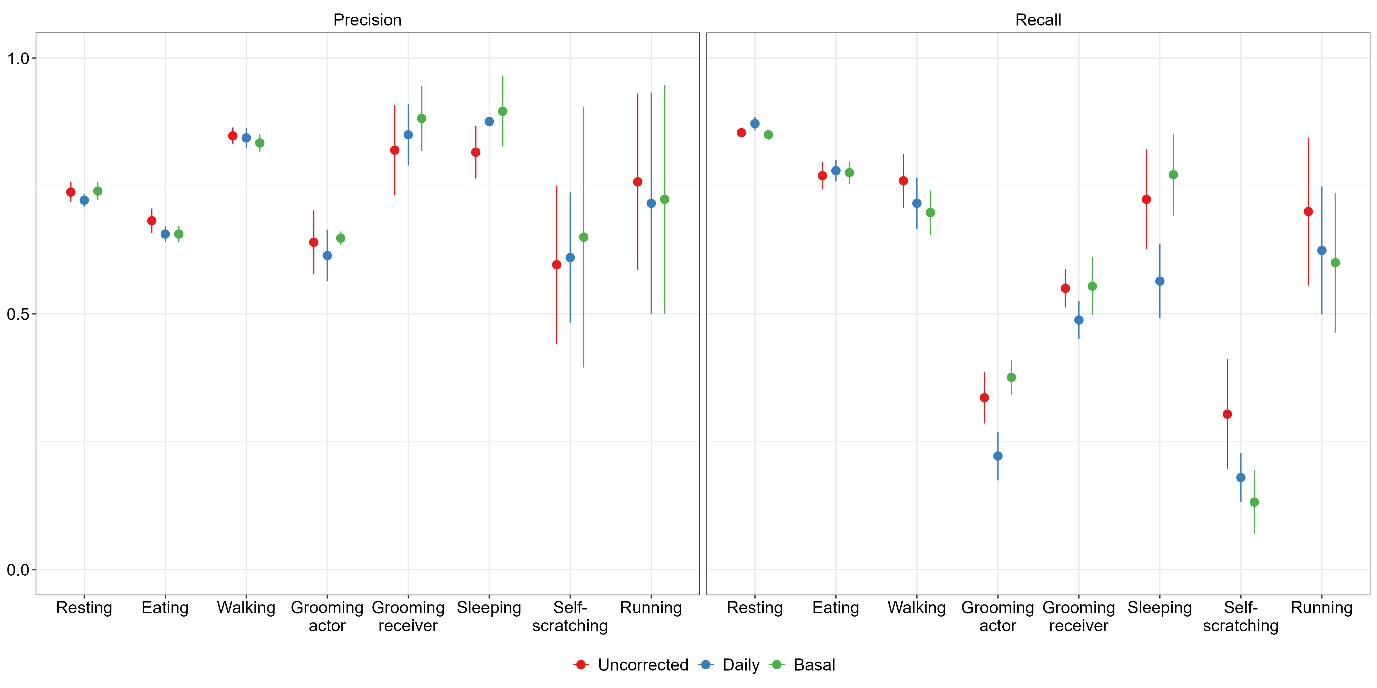

**Figure S5**: Mean precision and recall (± SD) across five training repeats of the TabPFN model using a burst length of 4 s under three orientation treatments (uncorrected, daily correction, and basal correction). Points represent means and error bars standard deviations. Differences among orientation treatments increase with behavioural rarity, with the largest divergence observed for uncommon and rare postural behaviours (e.g. grooming and sleeping), where basal correction yields higher recall.

##### Figure S6

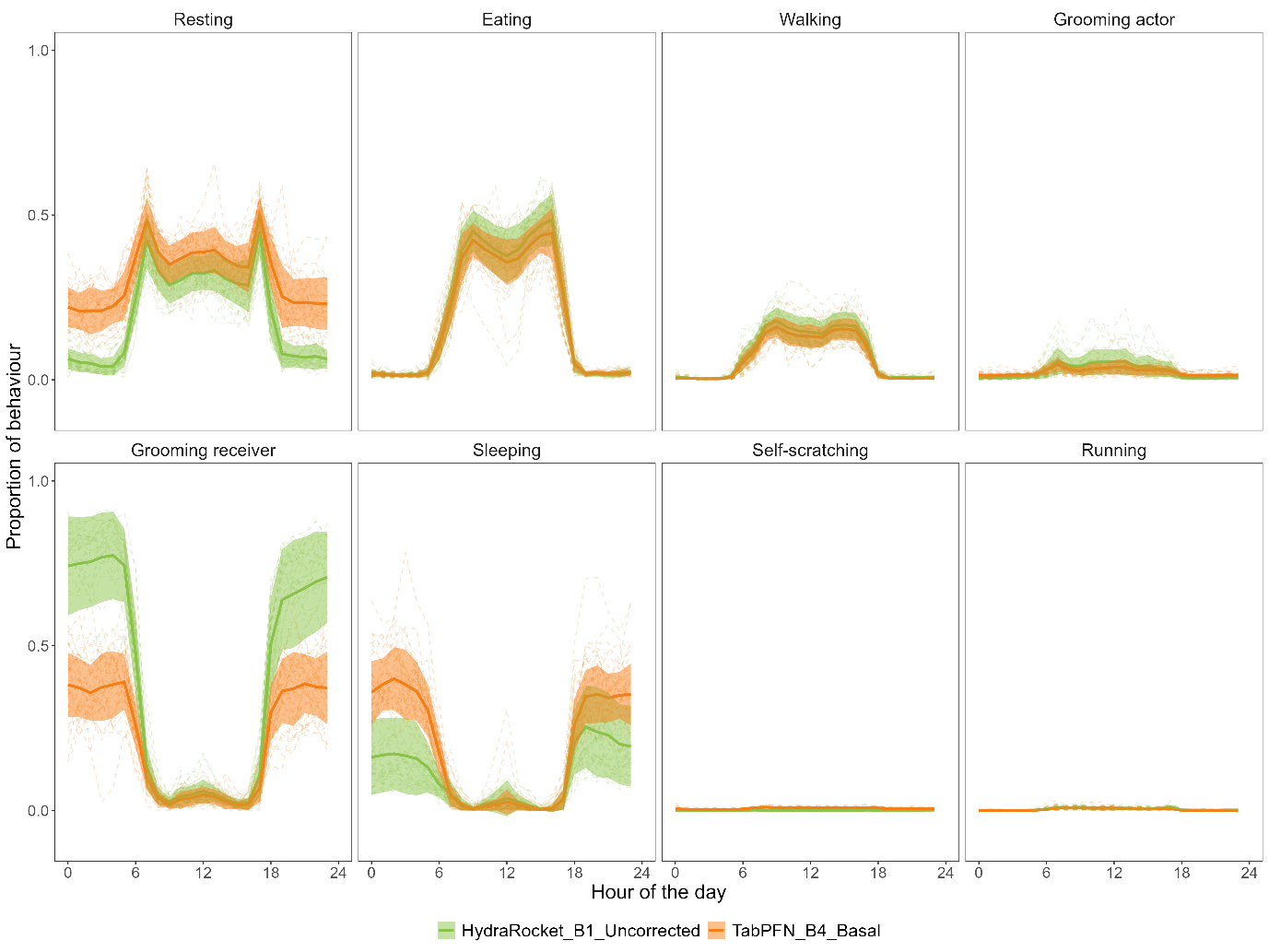

**Figure S6**: Circadian variation in predictions across behavioural classes from the two shortlisted models (HydraMultiROCKET trained on uncorrected burst length 1 dataset and TabPFN trained on basal corrected burst length 4 dataset) from seven randomly chosen days sampled within a three-month interval. Dotted lines represent data from different individuals (total n = 32) while the solid line and the shaded region around it represents the mean and standard deviation for the whole group. The lines are coloured based on the model the predictions are obtained from.

##### Figure S7

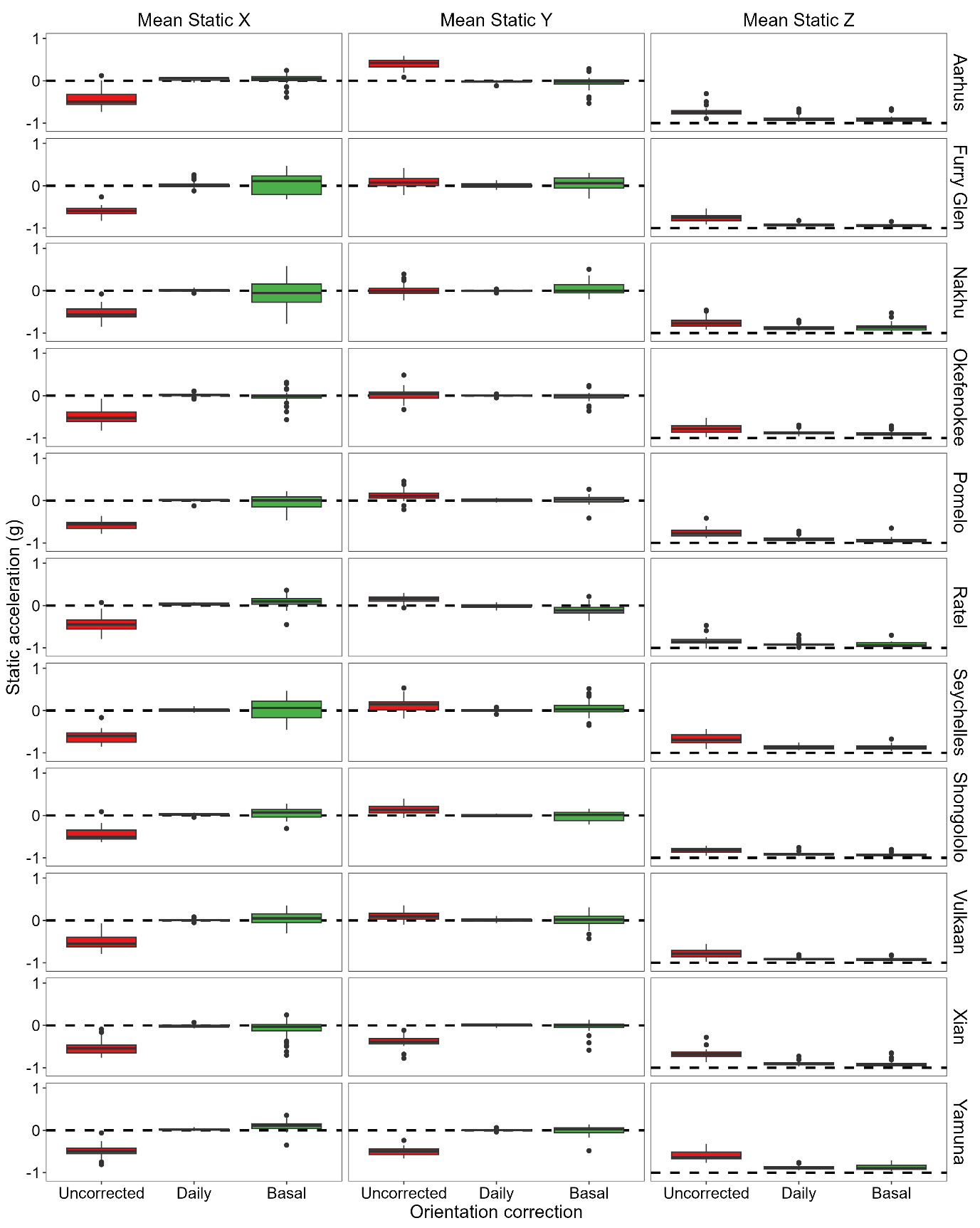

**Figure S7**: The mean static acceleration of X, Y and Z axes during bursts classified as resting from the raw uncorrected dataset and the daily and basal 3D orientation corrected datasets. Each facet represents data from one individual and one axis. Only data for individuals with at least 15 bursts assigned to resting are shown. Orientation correction defines a body-cantered reference frame based on walking bouts, with the Z-axis aligned to gravity and the horizontal axes defined relative to the estimated forward direction during locomotion. Under this reference frame, static acceleration during walking is expected to be close to 0 on the X and Y axes and −1 on the Z axis when alignment is stable. Deviations from these values during sleeping reflect differences between walking posture and sleeping posture, as well as residual variability in orientation estimation. Boxplots are coloured by the correction type.

### Supplementary Tables

##### Table S1:

Ethogram listing all behaviors used for video annotation, closely aligned with the focal-sampling protocol. Individual annotated behaviors were subsequently merged into a reduced set of *Final categories* (indicated in the corresponding column) to enable identification from accelerometer data. These final categories form the basis for burst-level behavioural attribution. Percentages indicate the proportion of bursts assigned to each final behavioural category for each burst length.

| **Behaviors** | **Definition** | **Final Category** | **Proportion by burst length [%]** | | | |
| --- | --- | --- | --- | --- | --- | --- |
|  |  |  | **1** | **2** | **3** | **4** |
| **Auto grooming** | Self-grooming | **Resting** | **38.04** | **36.95** | **36.64** | **36.32** |
| **Lying (resting)** | Lying down in any postures (dorsal, ventral or on the side) |  |  |  |  |  |
| **Sitting (resting)** | Sat motionless without being vigilant |  |  |  |  |  |
| **Standing** | Stationary in quadrupedal posture |  |  |  |  |  |
| **Chewing** | Individuals are sitting and chewing between foraging bouts |  |  |  |  |  |
| **Sitting (vigilant)** | Sitting while actively looking around, typically at the top of a tree |  |  |  |  |  |
| **Standing (vigilant)** | Stationary in quadrupedal posture while being vigilant |  |  |  |  |  |
| **Eating** | The action of handling something and consuming it | **Eating** | **28.83** | **28.10** | **27.80** | **27.51** |
| **Foraging (canopy)** | Actively searching for or consuming food while being in the trees |  |  |  |  |  |
| **Foraging (ground)** | Actively searching for or consuming food while being on the ground |  |  |  |  |  |
| **Canopy movement** | Any locomotion movement while being and staying in the trees | **Walking** | **10.59** | **10.77** | **10.65** | **11.24** |
| **Climbing downward** | Actively going down a tree |  |  |  |  |  |
| **Climbing upward** | Actively going up a tree |  |  |  |  |  |
| **Walking** | Moving on the ground with a walking gait |  |  |  |  |  |
| **Grooming (actor)** | Picking at or looking through fur of conspecifics | **Grooming (actor)** | **8.18** | **8.30** | **8.08** | **8.13** |
| **Grooming (receiver)** | Being groomed by a conspecific | **Grooming (receiver)** | **6.11** | **6.09** | **6.07** | **6.06** |
| **Sleeping** | Individuals stay motionless and with closed eyes in a lying or sitting position | **Sleeping** | **2.24** | **2.26** | **2.26** | **2.26** |
| **Self-Scratching** | Self-scratching with front or hind legs | **Self-Scratching** | **0.69** | **2.43** | **2.94** | **2.94** |
| **Running** | Moving on the ground with galloping gait | **Running** | **0.43** | **0.43** | **0.62** | **0.68** |
| **Chase** | Fleeing or chasing another individual |  |  |  |  |  |
| **Fighting** | Two individuals hit or bite each other, often accompanied by rapid movements such as jumping or running toward each other |  |  |  |  |  |
| **Jumping** | Act of jumping | **Discarded** | **4.91** | **4.68** | **4.94** | **4.80** |
| **Other complex locomotion** | Pivoting/spinning while walking |  |  |  |  |  |
| **River crossing** | Individuals are crossing the river |  |  |  |  |  |
| **Chest Rubbing** | A male displaying in front of another male by rubbing his chest against, most of the time a trunk |  |  |  |  |  |
| **Inspecting genitals** | Touching and/or inspecting an individual's genitals |  |  |  |  |  |
| **Masturbating** | When an individual engages in a sexual behaviour alone |  |  |  |  |  |
| **Muzzle contact** | Two individuals are touching each other's muzzles |  |  |  |  |  |
| **Playing** | When individuals are playing together |  |  |  |  |  |
| **Social** | Any other social interactions (embrace, lip smack, touching but not grooming, smell but not in a sexual way) |  |  |  |  |  |
| **Swimming** | Individual is swimming |  |  |  |  |  |
| **Vomiting** | Individuals regurgitate their food |  |  |  |  |  |
| **Body shake** | Fast whole-body movement from side to side |  |  |  |  |  |
| **Aggressive display** | Threatening body postures (stare, open mouth) |  |  |  |  |  |
| **Head Bob** | Individuals are moving their head up and down while staring at another in an agonistic context |  |  |  |  |  |
| **Mating** | Complete or partial mounting and/or copulating with an individual |  |  |  |  |  |
| **Standing up (vigilant)** | Stationary in bipedal posture, often looking around |  |  |  |  |  |
| **Threat** | Individuals detected a threat, are vocalizing, moving around, typically in presence of a predator or another group of monkeys |  |  |  |  |  |
| **Out of sight** | Subject is lost with unknown behavior | **Out of sight** | **-** | | | |

##### Table S2:

Overview of the machine-learning (ML) and deep-learning (DL) models evaluated in this study. Models are grouped by learning paradigm (ML or DL) and by whether they rely on engineered features or operate directly on raw time-series data. For each model, the table reports the model’s name, a brief description of its core principles and intended use, and the corresponding reference.

| DL or ML | Features based | Name | Description | Reference |
| --- | --- | --- | --- | --- |
| ML | yes | Support Vector Machine (SVM) | Linear margin-based classifier effective for separating non-linear patterns in feature space | Cortes & Vapnik, 1995 |
| ML | yes | Random Forest (RF) | Ensemble tree classifier widely used in accelerometer-based behavioural studies; robust to noisy and imbalanced data and acts as out baseline. | Breiman, 2001 |
| ML | yes | Extreme Gradient Boosting (XGBoost) | Gradient-boosted ensemble often outperforming RF on structured datasets. | Chen & Guestrin, 2016 |
| DL | yes | Category Embedding Model (CEM) | Neural network using embedding layers to learn interactions between engineered features. | Guo & Berkhahn, 2016 |
| DL | yes | Gated Adaptive Network for Deep Automated Learning of Features (GANDALF) | DL model for tabular data using gated mechanisms and built-in feature selection, designed to enhance learning in small datasets. | Joseph & Raj, 2022 |
| DL | yes | Tabular Prior-data Fitted Network (TabPFN) | Transformer-based model that performs tabular prediction via in-context learning. | Hollmann et al., 2025 |
| DL | no | Long Short-Term Memory (LSTM) | Recurrent neural network capturing temporal dependencies in sequential data; included as a baseline time-series model. | Hochreiter & Schmidhuber, 1997 |
| DL | no | Time-series Sequencer (TSSequencer) | 1D-convolutional sequence architecture integrating self-attention and recurrent modules to model raw time-series inputs. | Tatsunami & Taki, 2022 |
| DL | no | Hydra-MultiRocket (HMR) | State-of-the-art time-series classifier generating large convolutional feature banks; performs well on short and noisy windows. | Dempster, 2023 |

##### Table S3:

Summary of the engineered features extracted from accelerometer data and used as inputs for feature-based machine-learning and deep-learning models. Features include descriptive statistics per axis (x3) or burst (x1), orientation metrics, dynamic body acceleration measures, spectral characteristics, and principal-component–based descriptors capturing dominant patterns of dynamic acceleration variability. For each feature group, the number of derived variables is reported, yielding a total of 60 features per burst.

| **Features description** | **Number of features** |
| --- | --- |
| Mean, Standard deviation, Minimum, Maximum, Variance, Inverse Coefficient of Variation, Skewness, Kurtosis and Range (x,y,z) | 9x3 = 27 |
| Mean of pitch and roll | 2x1= 2 |
| Mean of static components (x,y,z) | 1x3 = 3 |
| Mean of Partial Dynamic Body Acceleration (PDBA) (x, y, z) | 1x3 = 3 |
| Mean of Overall Dynamic Body Acceleration (ODBA) | 1x1 =1 |
| Mean of Vectorized Dynamic Body Acceleration (VeDBA) | 1x1 =1 |
| Mean of smoothed VeDBA | 1x1 =1 |
| *q^2^* statistic (the vectorial sum of the three raw acceleration axes) | 1x1 =1 |
| Power spectral density (amplitude) and corresponding frequency of the first and second dominant spectral peaks (x, y, z) | 2x3x3 = 12 |
| Mean and variance of the first three PCA derived from PDBA values | 2x3x1=6 |
| Ratios of variance explained between the first three PCs (PC1/PC2; PC1/PC3; PC2/PC3) | 3x1=3 |
| **Total** | **60** |

##### Table S4:

The marginal means and 95% confidence intervals (CI_low_ – CI_high_) of accuracy and ROC AUC of all models compared in experiments 1-3.

| **Experiment** | **Model** | **Burst** | **Correction** | **Metric** | **Value** | **CI_low_** | **CI_high_** |
| --- | --- | --- | --- | --- | --- | --- | --- |
| 1 | RF | 1 | uncorrected | accuracy | 0.73 | 0.71 | 0.75 |
| 1 | RF | 2 | uncorrected | accuracy | 0.72 | 0.70 | 0.73 |
| 1 | RF | 3 | uncorrected | accuracy | 0.74 | 0.72 | 0.75 |
| 1 | RF | 4 | uncorrected | accuracy | 0.73 | 0.71 | 0.74 |
| 1 | RF | 1 | uncorrected | roc_auc | 0.92 | 0.91 | 0.92 |
| 1 | RF | 2 | uncorrected | roc_auc | 0.92 | 0.91 | 0.92 |
| 1 | RF | 3 | uncorrected | roc_auc | 0.92 | 0.91 | 0.93 |
| 1 | RF | 4 | uncorrected | roc_auc | 0.91 | 0.91 | 0.92 |
| 2 | RF | 1 | uncorrected | accuracy | 0.73 | 0.71 | 0.75 |
| 2 | RF | 1 | daily | accuracy | 0.69 | 0.67 | 0.71 |
| 2 | RF | 1 | basal | accuracy | 0.69 | 0.67 | 0.72 |
| 2 | RF | 1 | uncorrected | roc_auc | 0.92 | 0.91 | 0.92 |
| 2 | RF | 1 | daily | roc_auc | 0.90 | 0.89 | 0.91 |
| 2 | RF | 1 | basal | roc_auc | 0.90 | 0.89 | 0.91 |
| 3 | RF | 1 | uncorrected | accuracy | 0.73 | 0.70 | 0.76 |
| 3 | SVC | 1 | uncorrected | accuracy | 0.57 | 0.54 | 0.60 |
| 3 | XGB | 1 | uncorrected | accuracy | 0.72 | 0.70 | 0.75 |
| 3 | CatEmbed | 1 | uncorrected | accuracy | 0.72 | 0.69 | 0.74 |
| 3 | GANDALF | 1 | uncorrected | accuracy | 0.69 | 0.66 | 0.71 |
| 3 | TabPFN | 1 | uncorrected | accuracy | 0.76 | 0.74 | 0.79 |
| 3 | LSTM | 1 | uncorrected | accuracy | 0.40 | 0.37 | 0.43 |
| 3 | TSSeq | 1 | uncorrected | accuracy | 0.66 | 0.64 | 0.69 |
| 3 | HydraMultiROCKET | 1 | uncorrected | accuracy | 0.77 | 0.74 | 0.79 |
| 3 | RF | 1 | uncorrected | roc_auc | 0.91 | 0.90 | 0.93 |
| 3 | SVC | 1 | uncorrected | roc_auc | 0.84 | 0.82 | 0.86 |
| 3 | XGB | 1 | uncorrected | roc_auc | 0.91 | 0.90 | 0.93 |
| 3 | CatEmbed | 1 | uncorrected | roc_auc | 0.91 | 0.89 | 0.92 |
| 3 | GANDALF | 1 | uncorrected | roc_auc | 0.91 | 0.89 | 0.92 |
| 3 | TabPFN | 1 | uncorrected | roc_auc | 0.94 | 0.92 | 0.95 |
| 3 | LSTM | 1 | uncorrected | roc_auc | 0.61 | 0.59 | 0.64 |
| 3 | TSSeq | 1 | uncorrected | roc_auc | 0.86 | 0.84 | 0.88 |
| 3 | HydraMultiROCKET | 1 | uncorrected | roc_auc | 0.95 | 0.94 | 0.96 |

##### Table S5:

The difference, z value and p value of post-hoc comparisons between various levels of the main predictors from the statistical comparisons for experiments 1-3.

| **Experiment** | **Model** | **Burst** | **Correction** | **Metric** | **Level1** | **Level2** | **Difference** | **z** | **p** |
| --- | --- | --- | --- | --- | --- | --- | --- | --- | --- |
| 1 | RF |  | uncorrected | accuracy | Burst_2 | Burst_1 | -0.01 | -1.08 | 0.281 |
| 1 | RF |  | uncorrected | accuracy | Burst_3 | Burst_1 | 0.01 | 0.58 | 0.565 |
| 1 | RF |  | uncorrected | accuracy | Burst_4 | Burst_1 | 0.00 | -0.36 | 0.716 |
| 1 | RF |  | uncorrected | accuracy | Burst_3 | Burst_2 | 0.02 | 1.65 | 0.098 |
| 1 | RF |  | uncorrected | accuracy | Burst_4 | Burst_2 | 0.01 | 0.72 | 0.474 |
| 1 | RF |  | uncorrected | accuracy | Burst_4 | Burst_3 | -0.01 | -0.94 | 0.348 |
| 1 | RF |  | uncorrected | roc_auc | Burst_2 | Burst_1 | 0.00 | 0.04 | 0.972 |
| 1 | RF |  | uncorrected | roc_auc | Burst_3 | Burst_1 | 0.00 | 0.59 | 0.558 |
| 1 | RF |  | uncorrected | roc_auc | Burst_4 | Burst_1 | 0.00 | -0.45 | 0.655 |
| 1 | RF |  | uncorrected | roc_auc | Burst_3 | Burst_2 | 0.00 | 0.55 | 0.582 |
| 1 | RF |  | uncorrected | roc_auc | Burst_4 | Burst_2 | 0.00 | -0.48 | 0.630 |
| 1 | RF |  | uncorrected | roc_auc | Burst_4 | Burst_3 | -0.01 | -1.03 | 0.302 |
| 2 | RF | 1 |  | accuracy | daily | uncorrected | -0.04 | -2.52 | 0.012 |
| 2 | RF | 1 |  | accuracy | basal | uncorrected | -0.04 | -2.48 | 0.013 |
| 2 | RF | 1 |  | accuracy | basal | daily | 0.00 | 0.04 | 0.968 |
| 2 | RF | 1 |  | roc_auc | daily | uncorrected | -0.02 | -2.60 | 0.009 |
| 2 | RF | 1 |  | roc_auc | basal | uncorrected | -0.02 | -2.60 | 0.009 |
| 2 | RF | 1 |  | roc_auc | basal | daily | 0.00 | 0.00 | 0.999 |
| 3 |  | 1 | uncorrected | accuracy | SVC | RF | -0.16 | -8.08 | <0.001 |
| 3 |  | 1 | uncorrected | accuracy | XGB | RF | -0.01 | -0.45 | 0.653 |
| 3 |  | 1 | uncorrected | accuracy | CatEmbed | RF | -0.02 | -0.80 | 0.426 |
| 3 |  | 1 | uncorrected | accuracy | GANDALF | RF | -0.04 | -2.21 | 0.027 |
| 3 |  | 1 | uncorrected | accuracy | TabPFN | RF | 0.03 | 1.84 | 0.066 |
| 3 |  | 1 | uncorrected | accuracy | LSTM | RF | -0.33 | -16.65 | <0.001 |
| 3 |  | 1 | uncorrected | accuracy | TSSeq | RF | -0.07 | -3.37 | 0.001 |
| 3 |  | 1 | uncorrected | accuracy | HydraMultiROCKET | RF | 0.04 | 2.12 | 0.034 |
| 3 |  | 1 | uncorrected | accuracy | XGB | SVC | 0.15 | 7.62 | <0.001 |
| 3 |  | 1 | uncorrected | accuracy | CatEmbed | SVC | 0.15 | 7.27 | <0.001 |
| 3 |  | 1 | uncorrected | accuracy | GANDALF | SVC | 0.12 | 5.84 | <0.001 |
| 3 |  | 1 | uncorrected | accuracy | TabPFN | SVC | 0.20 | 9.97 | <0.001 |
| 3 |  | 1 | uncorrected | accuracy | LSTM | SVC | -0.17 | -8.08 | <0.001 |
| 3 |  | 1 | uncorrected | accuracy | TSSeq | SVC | 0.10 | 4.66 | <0.001 |
| 3 |  | 1 | uncorrected | accuracy | HydraMultiROCKET | SVC | 0.20 | 10.26 | <0.001 |
| 3 |  | 1 | uncorrected | accuracy | CatEmbed | XGB | -0.01 | -0.35 | 0.730 |
| 3 |  | 1 | uncorrected | accuracy | GANDALF | XGB | -0.03 | -1.76 | 0.079 |
| 3 |  | 1 | uncorrected | accuracy | TabPFN | XGB | 0.04 | 2.29 | 0.022 |
| 3 |  | 1 | uncorrected | accuracy | LSTM | XGB | -0.32 | -16.15 | <0.001 |
| 3 |  | 1 | uncorrected | accuracy | TSSeq | XGB | -0.06 | -2.92 | 0.003 |
| 3 |  | 1 | uncorrected | accuracy | HydraMultiROCKET | XGB | 0.05 | 2.57 | 0.010 |
| 3 |  | 1 | uncorrected | accuracy | GANDALF | CatEmbed | -0.03 | -1.41 | 0.159 |
| 3 |  | 1 | uncorrected | accuracy | TabPFN | CatEmbed | 0.05 | 2.64 | 0.008 |
| 3 |  | 1 | uncorrected | accuracy | LSTM | CatEmbed | -0.32 | -15.77 | <0.001 |
| 3 |  | 1 | uncorrected | accuracy | TSSeq | CatEmbed | -0.05 | -2.57 | 0.010 |
| 3 |  | 1 | uncorrected | accuracy | HydraMultiROCKET | CatEmbed | 0.05 | 2.92 | 0.003 |
| 3 |  | 1 | uncorrected | accuracy | TabPFN | GANDALF | 0.08 | 4.05 | <0.001 |
| 3 |  | 1 | uncorrected | accuracy | LSTM | GANDALF | -0.29 | -14.22 | <0.001 |
| 3 |  | 1 | uncorrected | accuracy | TSSeq | GANDALF | -0.02 | -1.16 | 0.245 |
| 3 |  | 1 | uncorrected | accuracy | HydraMultiROCKET | GANDALF | 0.08 | 4.34 | <0.001 |
| 3 |  | 1 | uncorrected | accuracy | LSTM | TabPFN | -0.37 | -18.73 | <0.001 |
| 3 |  | 1 | uncorrected | accuracy | TSSeq | TabPFN | -0.10 | -5.22 | <0.001 |
| 3 |  | 1 | uncorrected | accuracy | HydraMultiROCKET | TabPFN | 0.01 | 0.28 | 0.777 |
| 3 |  | 1 | uncorrected | accuracy | TSSeq | LSTM | 0.27 | 12.97 | <0.001 |
| 3 |  | 1 | uncorrected | accuracy | HydraMultiROCKET | LSTM | 0.37 | 19.05 | <0.001 |
| 3 |  | 1 | uncorrected | accuracy | HydraMultiROCKET | TSSeq | 0.10 | 5.51 | <0.001 |
| 3 |  | 1 | uncorrected | roc_auc | SVC | RF | -0.08 | -6.44 | <0.001 |
| 3 |  | 1 | uncorrected | roc_auc | XGB | RF | 0.00 | -0.25 | 0.801 |
| 3 |  | 1 | uncorrected | roc_auc | CatEmbed | RF | -0.01 | -0.76 | 0.446 |
| 3 |  | 1 | uncorrected | roc_auc | GANDALF | RF | -0.01 | -0.85 | 0.396 |
| 3 |  | 1 | uncorrected | roc_auc | TabPFN | RF | 0.02 | 2.17 | 0.030 |
| 3 |  | 1 | uncorrected | roc_auc | LSTM | RF | -0.30 | -20.80 | <0.001 |
| 3 |  | 1 | uncorrected | roc_auc | TSSeq | RF | -0.05 | -4.77 | <0.001 |
| 3 |  | 1 | uncorrected | roc_auc | HydraMultiROCKET | RF | 0.03 | 3.78 | 0.000 |
| 3 |  | 1 | uncorrected | roc_auc | XGB | SVC | 0.07 | 6.19 | <0.001 |
| 3 |  | 1 | uncorrected | roc_auc | CatEmbed | SVC | 0.07 | 5.69 | <0.001 |
| 3 |  | 1 | uncorrected | roc_auc | GANDALF | SVC | 0.07 | 5.60 | <0.001 |
| 3 |  | 1 | uncorrected | roc_auc | TabPFN | SVC | 0.10 | 8.54 | <0.001 |
| 3 |  | 1 | uncorrected | roc_auc | LSTM | SVC | -0.22 | -14.25 | <0.001 |
| 3 |  | 1 | uncorrected | roc_auc | TSSeq | SVC | 0.02 | 1.69 | 0.092 |
| 3 |  | 1 | uncorrected | roc_auc | HydraMultiROCKET | SVC | 0.11 | 10.07 | <0.001 |
| 3 |  | 1 | uncorrected | roc_auc | CatEmbed | XGB | -0.01 | -0.51 | 0.610 |
| 3 |  | 1 | uncorrected | roc_auc | GANDALF | XGB | -0.01 | -0.60 | 0.551 |
| 3 |  | 1 | uncorrected | roc_auc | TabPFN | XGB | 0.02 | 2.42 | 0.016 |
| 3 |  | 1 | uncorrected | roc_auc | LSTM | XGB | -0.30 | -20.55 | <0.001 |
| 3 |  | 1 | uncorrected | roc_auc | TSSeq | XGB | -0.05 | -4.52 | <0.001 |
| 3 |  | 1 | uncorrected | roc_auc | HydraMultiROCKET | XGB | 0.04 | 4.02 | <0.001 |
| 3 |  | 1 | uncorrected | roc_auc | GANDALF | CatEmbed | 0.00 | -0.09 | 0.931 |
| 3 |  | 1 | uncorrected | roc_auc | TabPFN | CatEmbed | 0.03 | 2.92 | 0.003 |
| 3 |  | 1 | uncorrected | roc_auc | LSTM | CatEmbed | -0.29 | -20.04 | <0.001 |
| 3 |  | 1 | uncorrected | roc_auc | TSSeq | CatEmbed | -0.05 | -4.01 | <0.001 |
| 3 |  | 1 | uncorrected | roc_auc | HydraMultiROCKET | CatEmbed | 0.04 | 4.53 | <0.001 |
| 3 |  | 1 | uncorrected | roc_auc | TabPFN | GANDALF | 0.03 | 3.01 | 0.003 |
| 3 |  | 1 | uncorrected | roc_auc | LSTM | GANDALF | -0.29 | -19.96 | <0.001 |
| 3 |  | 1 | uncorrected | roc_auc | TSSeq | GANDALF | -0.05 | -3.93 | <0.001 |
| 3 |  | 1 | uncorrected | roc_auc | HydraMultiROCKET | GANDALF | 0.04 | 4.61 | <0.001 |
| 3 |  | 1 | uncorrected | roc_auc | LSTM | TabPFN | -0.32 | -22.88 | <0.001 |
| 3 |  | 1 | uncorrected | roc_auc | TSSeq | TabPFN | -0.08 | -6.89 | <0.001 |
| 3 |  | 1 | uncorrected | roc_auc | HydraMultiROCKET | TabPFN | 0.01 | 1.63 | 0.104 |
| 3 |  | 1 | uncorrected | roc_auc | TSSeq | LSTM | 0.25 | 15.98 | <0.001 |
| 3 |  | 1 | uncorrected | roc_auc | HydraMultiROCKET | LSTM | 0.33 | 24.36 | <0.001 |
| 3 |  | 1 | uncorrected | roc_auc | HydraMultiROCKET | TSSeq | 0.09 | 8.44 | <0.001 |

##### Table S6:

The marginal means and 95% confidence intervals (CI_low_ – CI_high_) of accuracy and ROC AUC of the behavioural categories from all models compared in experiments 1-3.

| **Experiment** | **Model** | **Burst** | **Correction** | **Behavioral Category** | **Metric** | **Value** | **CI_low_** | **CI_high_** |
| --- | --- | --- | --- | --- | --- | --- | --- | --- |
| 1 | RF | 1 | uncorrected | common | precision | 0.72 | 0.66 | 0.78 |
| 1 | RF | 1 | uncorrected | uncommon | precision | 0.77 | 0.73 | 0.82 |
| 1 | RF | 2 | uncorrected | common | precision | 0.67 | 0.61 | 0.73 |
| 1 | RF | 2 | uncorrected | uncommon | precision | 0.73 | 0.68 | 0.79 |
| 1 | RF | 3 | uncorrected | common | precision | 0.71 | 0.65 | 0.77 |
| 1 | RF | 3 | uncorrected | uncommon | precision | 0.77 | 0.72 | 0.82 |
| 1 | RF | 4 | uncorrected | common | precision | 0.72 | 0.66 | 0.78 |
| 1 | RF | 4 | uncorrected | uncommon | precision | 0.78 | 0.73 | 0.82 |
| 1 | RF | 1 | uncorrected | common | recall | 0.79 | 0.73 | 0.84 |
| 1 | RF | 1 | uncorrected | uncommon | recall | 0.51 | 0.44 | 0.57 |
| 1 | RF | 2 | uncorrected | common | recall | 0.80 | 0.74 | 0.85 |
| 1 | RF | 2 | uncorrected | uncommon | recall | 0.52 | 0.45 | 0.59 |
| 1 | RF | 3 | uncorrected | common | recall | 0.81 | 0.76 | 0.86 |
| 1 | RF | 3 | uncorrected | uncommon | recall | 0.54 | 0.47 | 0.61 |
| 1 | RF | 4 | uncorrected | common | recall | 0.80 | 0.75 | 0.85 |
| 1 | RF | 4 | uncorrected | uncommon | recall | 0.52 | 0.45 | 0.59 |
| 2 | RF | 4 | uncorrected | common | precision | 0.71 | 0.61 | 0.82 |
| 2 | RF | 4 | uncorrected | uncommon | precision | 0.79 | 0.71 | 0.86 |
| 2 | RF | 4 | uncorrected | rare | precision | 0.80 | 0.74 | 0.87 |
| 2 | RF | 4 | daily | common | precision | 0.57 | 0.46 | 0.69 |
| 2 | RF | 4 | daily | uncommon | precision | 0.67 | 0.57 | 0.76 |
| 2 | RF | 4 | daily | rare | precision | 0.69 | 0.60 | 0.79 |
| 2 | RF | 4 | basal | common | precision | 0.62 | 0.51 | 0.74 |
| 2 | RF | 4 | basal | uncommon | precision | 0.71 | 0.62 | 0.80 |
| 2 | RF | 4 | basal | rare | precision | 0.73 | 0.65 | 0.82 |
| 2 | RF | 4 | uncorrected | common | recall | 0.80 | 0.74 | 0.87 |
| 2 | RF | 4 | uncorrected | uncommon | recall | 0.55 | 0.47 | 0.64 |
| 2 | RF | 4 | uncorrected | rare | recall | 0.40 | 0.32 | 0.49 |
| 2 | RF | 4 | daily | common | recall | 0.73 | 0.65 | 0.81 |
| 2 | RF | 4 | daily | uncommon | recall | 0.45 | 0.37 | 0.53 |
| 2 | RF | 4 | daily | rare | recall | 0.31 | 0.24 | 0.38 |
| 2 | RF | 4 | basal | common | recall | 0.74 | 0.67 | 0.82 |
| 2 | RF | 4 | basal | uncommon | recall | 0.47 | 0.38 | 0.55 |
| 2 | RF | 4 | basal | rare | recall | 0.32 | 0.25 | 0.40 |
| 3 | RF | 4 | uncorrected | common | precision | 0.65 | 0.51 | 0.79 |
| 3 | RF | 4 | uncorrected | uncommon | precision | 0.73 | 0.63 | 0.83 |
| 3 | RF | 4 | uncorrected | rare | precision | 0.88 | 0.83 | 0.94 |
| 3 | SVC | 4 | uncorrected | common | precision | 0.57 | 0.42 | 0.71 |
| 3 | SVC | 4 | uncorrected | uncommon | precision | 0.31 | 0.20 | 0.41 |
| 3 | SVC | 4 | uncorrected | rare | precision | 0.11 | 0.06 | 0.16 |
| 3 | XGB | 4 | uncorrected | common | precision | 0.65 | 0.51 | 0.79 |
| 3 | XGB | 4 | uncorrected | uncommon | precision | 0.71 | 0.60 | 0.81 |
| 3 | XGB | 4 | uncorrected | rare | precision | 0.73 | 0.63 | 0.83 |
| 3 | CatEmbed | 4 | uncorrected | common | precision | 0.65 | 0.51 | 0.79 |
| 3 | CatEmbed | 4 | uncorrected | uncommon | precision | 0.69 | 0.58 | 0.79 |
| 3 | CatEmbed | 4 | uncorrected | rare | precision | 0.52 | 0.40 | 0.64 |
| 3 | GANDALF | 4 | uncorrected | common | precision | 0.64 | 0.50 | 0.78 |
| 3 | GANDALF | 4 | uncorrected | uncommon | precision | 0.66 | 0.55 | 0.77 |
| 3 | GANDALF | 4 | uncorrected | rare | precision | 0.52 | 0.40 | 0.64 |
| 3 | TabPFN | 4 | uncorrected | common | precision | 0.65 | 0.52 | 0.79 |
| 3 | TabPFN | 4 | uncorrected | uncommon | precision | 0.71 | 0.61 | 0.82 |
| 3 | TabPFN | 4 | uncorrected | rare | precision | 0.71 | 0.61 | 0.82 |
| 3 | LSTM | 4 | uncorrected | common | precision | 0.61 | 0.47 | 0.76 |
| 3 | LSTM | 4 | uncorrected | uncommon | precision | 0.46 | 0.34 | 0.58 |
| 3 | LSTM | 4 | uncorrected | rare | precision | 0.21 | 0.12 | 0.29 |
| 3 | TSSeq | 4 | uncorrected | common | precision | 0.64 | 0.50 | 0.78 |
| 3 | TSSeq | 4 | uncorrected | uncommon | precision | 0.65 | 0.54 | 0.77 |
| 3 | TSSeq | 4 | uncorrected | rare | precision | 0.65 | 0.54 | 0.76 |
| 3 | HydraMultiROCKET | 4 | uncorrected | common | precision | 0.67 | 0.53 | 0.80 |
| 3 | HydraMultiROCKET | 4 | uncorrected | uncommon | precision | 0.68 | 0.57 | 0.78 |
| 3 | HydraMultiROCKET | 4 | uncorrected | rare | precision | 0.79 | 0.70 | 0.87 |
| 3 | RF | 4 | uncorrected | common | recall | 0.76 | 0.66 | 0.87 |
| 3 | RF | 4 | uncorrected | uncommon | recall | 0.52 | 0.41 | 0.63 |
| 3 | RF | 4 | uncorrected | rare | recall | 0.47 | 0.36 | 0.58 |
| 3 | SVC | 4 | uncorrected | common | recall | 0.75 | 0.65 | 0.86 |
| 3 | SVC | 4 | uncorrected | uncommon | recall | 0.14 | 0.08 | 0.20 |
| 3 | SVC | 4 | uncorrected | rare | recall | 0.08 | 0.04 | 0.12 |
| 3 | XGB | 4 | uncorrected | common | recall | 0.75 | 0.64 | 0.86 |
| 3 | XGB | 4 | uncorrected | uncommon | recall | 0.54 | 0.43 | 0.65 |
| 3 | XGB | 4 | uncorrected | rare | recall | 0.50 | 0.39 | 0.61 |
| 3 | CatEmbed | 4 | uncorrected | common | recall | 0.74 | 0.63 | 0.85 |
| 3 | CatEmbed | 4 | uncorrected | uncommon | recall | 0.52 | 0.41 | 0.63 |
| 3 | CatEmbed | 4 | uncorrected | rare | recall | 0.41 | 0.30 | 0.51 |
| 3 | GANDALF | 4 | uncorrected | common | recall | 0.75 | 0.64 | 0.85 |
| 3 | GANDALF | 4 | uncorrected | uncommon | recall | 0.48 | 0.37 | 0.58 |
| 3 | GANDALF | 4 | uncorrected | rare | recall | 0.30 | 0.21 | 0.40 |
| 3 | TabPFN | 4 | uncorrected | common | recall | 0.75 | 0.65 | 0.86 |
| 3 | TabPFN | 4 | uncorrected | uncommon | recall | 0.54 | 0.44 | 0.65 |
| 3 | TabPFN | 4 | uncorrected | rare | recall | 0.57 | 0.46 | 0.68 |
| 3 | LSTM | 4 | uncorrected | common | recall | 0.73 | 0.62 | 0.84 |
| 3 | LSTM | 4 | uncorrected | uncommon | recall | 0.31 | 0.21 | 0.41 |
| 3 | LSTM | 4 | uncorrected | rare | recall | 0.17 | 0.10 | 0.24 |
| 3 | TSSeq | 4 | uncorrected | common | recall | 0.74 | 0.64 | 0.85 |
| 3 | TSSeq | 4 | uncorrected | uncommon | recall | 0.51 | 0.40 | 0.62 |
| 3 | TSSeq | 4 | uncorrected | rare | recall | 0.28 | 0.19 | 0.37 |
| 3 | HydraMultiROCKET | 4 | uncorrected | common | recall | 0.75 | 0.64 | 0.86 |
| 3 | HydraMultiROCKET | 4 | uncorrected | uncommon | recall | 0.57 | 0.46 | 0.67 |
| 3 | HydraMultiROCKET | 4 | uncorrected | rare | recall | 0.60 | 0.49 | 0.70 |

##### Table S7:

The difference, z value and p value of post-hoc comparisons across behavioural categories between various levels of the main predictors from the statistical comparisons for experiments 1-3.

| **Experiment** | **Model** | **Burst** | **Correction** | **metric** | **Level1** | **Level2** | **Difference** | **z** | **p** |
| --- | --- | --- | --- | --- | --- | --- | --- | --- | --- |
| 1 | RF |  | uncorrected | precision | uncommon | common | 0.06 | 2.83 | 0.005 |
| 1 | RF |  | uncorrected | recall | uncommon | common | -0.27 | -10.07 | < 0.001 |
| 2 | RF | 4 |  | precision | uncommon | common | 0.08 | 1.64 | 0.101 |
| 2 | RF | 4 |  | precision | rare | common | 0.11 | 2.13 | 0.033 |
| 2 | RF | 4 |  | precision | rare | uncommon | 0.02 | 0.56 | 0.579 |
| 2 | RF | 4 |  | recall | uncommon | common | -0.27 | -5.98 | < 0.001 |
| 2 | RF | 4 |  | recall | rare | common | -0.41 | -9.47 | < 0.001 |
| 2 | RF | 4 |  | recall | rare | uncommon | -0.15 | -3.37 | 0.001 |
| 3 |  | 4 | uncorrected | precision | uncommon | common | -0.01 | -0.47 | 0.639 |
| 3 |  | 4 | uncorrected | precision | rare | common | -0.07 | -2.33 | 0.020 |
| 3 |  | 4 | uncorrected | precision | rare | uncommon | -0.05 | -2.15 | 0.031 |
| 3 |  | 4 | uncorrected | recall | uncommon | common | -0.29 | -11.27 | < 0.001 |
| 3 |  | 4 | uncorrected | recall | rare | common | -0.37 | -14.95 | < 0.001 |
| 3 |  | 4 | uncorrected | recall | rare | uncommon | -0.08 | -3.53 | < 0.001 |

##### Table S8:

The difference, z value and p value of post-hoc comparisons within behavioural categories across various levels of the main predictors from the statistical comparisons for experiments 1-3. In the statistical analyses for experiments 1 and 2, there was no significant interaction between the behavioural categories and the main predictors, so we the contrast comparison is performed based on the average behavioural category value.

| **Experiment** | **Model** | **Burst** | **Correction** | **Behavioral category** | **Metric** | **Level1** | **Level2** | **Difference** | **z** | **p** |
| --- | --- | --- | --- | --- | --- | --- | --- | --- | --- | --- |
| 1 | RF |  | uncorrected |  | precision | Burst_2 | Burst_1 | -0.04 | -1.55 | 0.122 |
| 1 | RF |  | uncorrected |  | precision | Burst_3 | Burst_1 | -0.01 | -0.26 | 0.793 |
| 1 | RF |  | uncorrected |  | precision | Burst_4 | Burst_1 | 0.00 | 0.06 | 0.954 |
| 1 | RF |  | uncorrected |  | precision | Burst_3 | Burst_2 | 0.04 | 1.28 | 0.199 |
| 1 | RF |  | uncorrected |  | precision | Burst_4 | Burst_2 | 0.05 | 1.61 | 0.108 |
| 1 | RF |  | uncorrected |  | precision | Burst_4 | Burst_3 | 0.01 | 0.32 | 0.747 |
| 1 | RF |  | uncorrected |  | recall | Burst_2 | Burst_1 | 0.01 | 0.31 | 0.754 |
| 1 | RF |  | uncorrected |  | recall | Burst_3 | Burst_1 | 0.03 | 0.74 | 0.457 |
| 1 | RF |  | uncorrected |  | recall | Burst_4 | Burst_1 | 0.01 | 0.37 | 0.708 |
| 1 | RF |  | uncorrected |  | recall | Burst_3 | Burst_2 | 0.02 | 0.43 | 0.670 |
| 1 | RF |  | uncorrected |  | recall | Burst_4 | Burst_2 | 0.00 | 0.06 | 0.952 |
| 1 | RF |  | uncorrected |  | recall | Burst_4 | Burst_3 | -0.01 | -0.37 | 0.714 |
| 2 | RF | 4 |  |  | precision | daily | uncorrected | -0.12 | -2.75 | 0.006 |
| 2 | RF | 4 |  |  | precision | basal | uncorrected | -0.08 | -1.84 | 0.065 |
| 2 | RF | 4 |  |  | precision | basal | daily | 0.04 | 0.94 | 0.350 |
| 2 | RF | 4 |  |  | recall | daily | uncorrected | -0.09 | -2.11 | 0.035 |
| 2 | RF | 4 |  |  | recall | basal | uncorrected | -0.08 | -1.77 | 0.076 |
| 2 | RF | 4 |  |  | recall | basal | daily | 0.01 | 0.33 | 0.738 |
| 3 |  | 4 | uncorrected | common | precision | SVC | RF | -0.08 | -0.79 | 0.430 |
| 3 |  | 4 | uncorrected | common | precision | XGB | RF | 0.00 | 0.03 | 0.973 |
| 3 |  | 4 | uncorrected | common | precision | CatEmbed | RF | 0.00 | 0.03 | 0.975 |
| 3 |  | 4 | uncorrected | common | precision | GANDALF | RF | -0.01 | -0.10 | 0.918 |
| 3 |  | 4 | uncorrected | common | precision | TabPFN | RF | 0.01 | 0.06 | 0.950 |
| 3 |  | 4 | uncorrected | common | precision | LSTM | RF | -0.03 | -0.34 | 0.735 |
| 3 |  | 4 | uncorrected | common | precision | TSSeq | RF | -0.01 | -0.09 | 0.929 |
| 3 |  | 4 | uncorrected | common | precision | HydraMultiROCKET | RF | 0.02 | 0.20 | 0.841 |
| 3 |  | 4 | uncorrected | common | precision | XGB | SVC | 0.08 | 0.82 | 0.411 |
| 3 |  | 4 | uncorrected | common | precision | CatEmbed | SVC | 0.08 | 0.82 | 0.412 |
| 3 |  | 4 | uncorrected | common | precision | GANDALF | SVC | 0.07 | 0.68 | 0.494 |
| 3 |  | 4 | uncorrected | common | precision | TabPFN | SVC | 0.09 | 0.85 | 0.394 |
| 3 |  | 4 | uncorrected | common | precision | LSTM | SVC | 0.05 | 0.45 | 0.654 |
| 3 |  | 4 | uncorrected | common | precision | TSSeq | SVC | 0.07 | 0.70 | 0.485 |
| 3 |  | 4 | uncorrected | common | precision | HydraMultiROCKET | SVC | 0.10 | 0.99 | 0.322 |
| 3 |  | 4 | uncorrected | common | precision | CatEmbed | XGB | 0.00 | 0.00 | 0.998 |
| 3 |  | 4 | uncorrected | common | precision | GANDALF | XGB | -0.01 | -0.14 | 0.891 |
| 3 |  | 4 | uncorrected | common | precision | TabPFN | XGB | 0.00 | 0.03 | 0.977 |
| 3 |  | 4 | uncorrected | common | precision | LSTM | XGB | -0.04 | -0.37 | 0.710 |
| 3 |  | 4 | uncorrected | common | precision | TSSeq | XGB | -0.01 | -0.12 | 0.902 |
| 3 |  | 4 | uncorrected | common | precision | HydraMultiROCKET | XGB | 0.02 | 0.17 | 0.868 |
| 3 |  | 4 | uncorrected | common | precision | GANDALF | CatEmbed | -0.01 | -0.14 | 0.892 |
| 3 |  | 4 | uncorrected | common | precision | TabPFN | CatEmbed | 0.00 | 0.03 | 0.975 |
| 3 |  | 4 | uncorrected | common | precision | LSTM | CatEmbed | -0.04 | -0.37 | 0.711 |
| 3 |  | 4 | uncorrected | common | precision | TSSeq | CatEmbed | -0.01 | -0.12 | 0.904 |
| 3 |  | 4 | uncorrected | common | precision | HydraMultiROCKET | CatEmbed | 0.02 | 0.17 | 0.866 |
| 3 |  | 4 | uncorrected | common | precision | TabPFN | GANDALF | 0.02 | 0.17 | 0.867 |
| 3 |  | 4 | uncorrected | common | precision | LSTM | GANDALF | -0.02 | -0.23 | 0.814 |
| 3 |  | 4 | uncorrected | common | precision | TSSeq | GANDALF | 0.00 | 0.01 | 0.988 |
| 3 |  | 4 | uncorrected | common | precision | HydraMultiROCKET | GANDALF | 0.03 | 0.30 | 0.761 |
| 3 |  | 4 | uncorrected | common | precision | LSTM | TabPFN | -0.04 | -0.40 | 0.688 |
| 3 |  | 4 | uncorrected | common | precision | TSSeq | TabPFN | -0.02 | -0.15 | 0.879 |
| 3 |  | 4 | uncorrected | common | precision | HydraMultiROCKET | TabPFN | 0.01 | 0.14 | 0.891 |
| 3 |  | 4 | uncorrected | common | precision | TSSeq | LSTM | 0.03 | 0.25 | 0.803 |
| 3 |  | 4 | uncorrected | common | precision | HydraMultiROCKET | LSTM | 0.05 | 0.54 | 0.590 |
| 3 |  | 4 | uncorrected | common | precision | HydraMultiROCKET | TSSeq | 0.03 | 0.29 | 0.772 |
| 3 |  | 4 | uncorrected | uncommon | precision | SVC | RF | -0.42 | -5.71 | < 0.001 |
| 3 |  | 4 | uncorrected | uncommon | precision | XGB | RF | -0.03 | -0.38 | 0.706 |
| 3 |  | 4 | uncorrected | uncommon | precision | CatEmbed | RF | -0.05 | -0.62 | 0.535 |
| 3 |  | 4 | uncorrected | uncommon | precision | GANDALF | RF | -0.07 | -0.93 | 0.352 |
| 3 |  | 4 | uncorrected | uncommon | precision | TabPFN | RF | -0.02 | -0.26 | 0.794 |
| 3 |  | 4 | uncorrected | uncommon | precision | LSTM | RF | -0.27 | -3.42 | 0.001 |
| 3 |  | 4 | uncorrected | uncommon | precision | TSSeq | RF | -0.08 | -1.04 | 0.300 |
| 3 |  | 4 | uncorrected | uncommon | precision | HydraMultiROCKET | RF | -0.06 | -0.77 | 0.441 |
| 3 |  | 4 | uncorrected | uncommon | precision | XGB | SVC | 0.40 | 5.21 | < 0.001 |
| 3 |  | 4 | uncorrected | uncommon | precision | CatEmbed | SVC | 0.38 | 4.90 | < 0.001 |
| 3 |  | 4 | uncorrected | uncommon | precision | GANDALF | SVC | 0.35 | 4.51 | < 0.001 |
| 3 |  | 4 | uncorrected | uncommon | precision | TabPFN | SVC | 0.41 | 5.36 | < 0.001 |
| 3 |  | 4 | uncorrected | uncommon | precision | LSTM | SVC | 0.15 | 1.84 | 0.065 |
| 3 |  | 4 | uncorrected | uncommon | precision | TSSeq | SVC | 0.35 | 4.38 | < 0.001 |
| 3 |  | 4 | uncorrected | uncommon | precision | HydraMultiROCKET | SVC | 0.37 | 4.71 | < 0.001 |
| 3 |  | 4 | uncorrected | uncommon | precision | CatEmbed | XGB | -0.02 | -0.24 | 0.807 |
| 3 |  | 4 | uncorrected | uncommon | precision | GANDALF | XGB | -0.04 | -0.55 | 0.580 |
| 3 |  | 4 | uncorrected | uncommon | precision | TabPFN | XGB | 0.01 | 0.12 | 0.908 |
| 3 |  | 4 | uncorrected | uncommon | precision | LSTM | XGB | -0.25 | -3.01 | 0.003 |
| 3 |  | 4 | uncorrected | uncommon | precision | TSSeq | XGB | -0.05 | -0.66 | 0.510 |
| 3 |  | 4 | uncorrected | uncommon | precision | HydraMultiROCKET | XGB | -0.03 | -0.39 | 0.694 |
| 3 |  | 4 | uncorrected | uncommon | precision | GANDALF | CatEmbed | -0.02 | -0.31 | 0.757 |
| 3 |  | 4 | uncorrected | uncommon | precision | TabPFN | CatEmbed | 0.03 | 0.36 | 0.719 |
| 3 |  | 4 | uncorrected | uncommon | precision | LSTM | CatEmbed | -0.23 | -2.75 | 0.006 |
| 3 |  | 4 | uncorrected | uncommon | precision | TSSeq | CatEmbed | -0.03 | -0.41 | 0.679 |
| 3 |  | 4 | uncorrected | uncommon | precision | HydraMultiROCKET | CatEmbed | -0.01 | -0.15 | 0.881 |
| 3 |  | 4 | uncorrected | uncommon | precision | TabPFN | GANDALF | 0.05 | 0.67 | 0.503 |
| 3 |  | 4 | uncorrected | uncommon | precision | LSTM | GANDALF | -0.20 | -2.42 | 0.016 |
| 3 |  | 4 | uncorrected | uncommon | precision | TSSeq | GANDALF | -0.01 | -0.10 | 0.917 |
| 3 |  | 4 | uncorrected | uncommon | precision | HydraMultiROCKET | GANDALF | 0.01 | 0.16 | 0.873 |
| 3 |  | 4 | uncorrected | uncommon | precision | LSTM | TabPFN | -0.25 | -3.14 | 0.002 |
| 3 |  | 4 | uncorrected | uncommon | precision | TSSeq | TabPFN | -0.06 | -0.77 | 0.439 |
| 3 |  | 4 | uncorrected | uncommon | precision | HydraMultiROCKET | TabPFN | -0.04 | -0.51 | 0.611 |
| 3 |  | 4 | uncorrected | uncommon | precision | TSSeq | LSTM | 0.19 | 2.31 | 0.021 |
| 3 |  | 4 | uncorrected | uncommon | precision | HydraMultiROCKET | LSTM | 0.22 | 2.59 | 0.010 |
| 3 |  | 4 | uncorrected | uncommon | precision | HydraMultiROCKET | TSSeq | 0.02 | 0.26 | 0.792 |
| 3 |  | 4 | uncorrected | rare | precision | SVC | RF | -0.78 | -20.05 | < 0.001 |
| 3 |  | 4 | uncorrected | rare | precision | XGB | RF | -0.15 | -2.67 | 0.007 |
| 3 |  | 4 | uncorrected | rare | precision | CatEmbed | RF | -0.36 | -5.32 | < 0.001 |
| 3 |  | 4 | uncorrected | rare | precision | GANDALF | RF | -0.36 | -5.35 | < 0.001 |
| 3 |  | 4 | uncorrected | rare | precision | TabPFN | RF | -0.17 | -2.89 | 0.004 |
| 3 |  | 4 | uncorrected | rare | precision | LSTM | RF | -0.67 | -12.93 | < 0.001 |
| 3 |  | 4 | uncorrected | rare | precision | TSSeq | RF | -0.23 | -3.67 | 0.000 |
| 3 |  | 4 | uncorrected | rare | precision | HydraMultiROCKET | RF | -0.10 | -1.85 | 0.064 |
| 3 |  | 4 | uncorrected | rare | precision | XGB | SVC | 0.62 | 10.83 | < 0.001 |
| 3 |  | 4 | uncorrected | rare | precision | CatEmbed | SVC | 0.41 | 6.18 | < 0.001 |
| 3 |  | 4 | uncorrected | rare | precision | GANDALF | SVC | 0.41 | 6.15 | < 0.001 |
| 3 |  | 4 | uncorrected | rare | precision | TabPFN | SVC | 0.61 | 10.29 | < 0.001 |
| 3 |  | 4 | uncorrected | rare | precision | LSTM | SVC | 0.10 | 2.03 | 0.042 |
| 3 |  | 4 | uncorrected | rare | precision | TSSeq | SVC | 0.54 | 8.60 | < 0.001 |
| 3 |  | 4 | uncorrected | rare | precision | HydraMultiROCKET | SVC | 0.68 | 13.22 | < 0.001 |
| 3 |  | 4 | uncorrected | rare | precision | CatEmbed | XGB | -0.21 | -2.58 | 0.010 |
| 3 |  | 4 | uncorrected | rare | precision | GANDALF | XGB | -0.21 | -2.61 | 0.009 |
| 3 |  | 4 | uncorrected | rare | precision | TabPFN | XGB | -0.02 | -0.23 | 0.821 |
| 3 |  | 4 | uncorrected | rare | precision | LSTM | XGB | -0.52 | -7.73 | < 0.001 |
| 3 |  | 4 | uncorrected | rare | precision | TSSeq | XGB | -0.08 | -1.03 | 0.303 |
| 3 |  | 4 | uncorrected | rare | precision | HydraMultiROCKET | XGB | 0.06 | 0.88 | 0.381 |
| 3 |  | 4 | uncorrected | rare | precision | GANDALF | CatEmbed | 0.00 | -0.02 | 0.981 |
| 3 |  | 4 | uncorrected | rare | precision | TabPFN | CatEmbed | 0.19 | 2.35 | 0.019 |
| 3 |  | 4 | uncorrected | rare | precision | LSTM | CatEmbed | -0.31 | -4.14 | < 0.001 |
| 3 |  | 4 | uncorrected | rare | precision | TSSeq | CatEmbed | 0.13 | 1.52 | 0.129 |
| 3 |  | 4 | uncorrected | rare | precision | HydraMultiROCKET | CatEmbed | 0.27 | 3.50 | 0.000 |
| 3 |  | 4 | uncorrected | rare | precision | TabPFN | GANDALF | 0.19 | 2.37 | 0.018 |
| 3 |  | 4 | uncorrected | rare | precision | LSTM | GANDALF | -0.31 | -4.11 | < 0.001 |
| 3 |  | 4 | uncorrected | rare | precision | TSSeq | GANDALF | 0.13 | 1.54 | 0.123 |
| 3 |  | 4 | uncorrected | rare | precision | HydraMultiROCKET | GANDALF | 0.27 | 3.53 | 0.000 |
| 3 |  | 4 | uncorrected | rare | precision | LSTM | TabPFN | -0.50 | -7.36 | < 0.001 |
| 3 |  | 4 | uncorrected | rare | precision | TSSeq | TabPFN | -0.06 | -0.80 | 0.422 |
| 3 |  | 4 | uncorrected | rare | precision | HydraMultiROCKET | TabPFN | 0.07 | 1.10 | 0.271 |
| 3 |  | 4 | uncorrected | rare | precision | TSSeq | LSTM | 0.44 | 6.11 | < 0.001 |
| 3 |  | 4 | uncorrected | rare | precision | HydraMultiROCKET | LSTM | 0.58 | 9.30 | < 0.001 |
| 3 |  | 4 | uncorrected | rare | precision | HydraMultiROCKET | TSSeq | 0.14 | 1.91 | 0.057 |
| 3 |  | 4 | uncorrected | common | recall | SVC | RF | -0.01 | -0.14 | 0.886 |
| 3 |  | 4 | uncorrected | common | recall | XGB | RF | -0.01 | -0.18 | 0.858 |
| 3 |  | 4 | uncorrected | common | recall | CatEmbed | RF | -0.02 | -0.32 | 0.753 |
| 3 |  | 4 | uncorrected | common | recall | GANDALF | RF | -0.02 | -0.23 | 0.822 |
| 3 |  | 4 | uncorrected | common | recall | TabPFN | RF | -0.01 | -0.15 | 0.883 |
| 3 |  | 4 | uncorrected | common | recall | LSTM | RF | -0.03 | -0.39 | 0.693 |
| 3 |  | 4 | uncorrected | common | recall | TSSeq | RF | -0.02 | -0.24 | 0.808 |
| 3 |  | 4 | uncorrected | common | recall | HydraMultiROCKET | RF | -0.01 | -0.19 | 0.846 |
| 3 |  | 4 | uncorrected | common | recall | XGB | SVC | 0.00 | -0.04 | 0.972 |
| 3 |  | 4 | uncorrected | common | recall | CatEmbed | SVC | -0.01 | -0.17 | 0.863 |
| 3 |  | 4 | uncorrected | common | recall | GANDALF | SVC | -0.01 | -0.08 | 0.934 |
| 3 |  | 4 | uncorrected | common | recall | TabPFN | SVC | 0.00 | 0.00 | 0.997 |
| 3 |  | 4 | uncorrected | common | recall | LSTM | SVC | -0.02 | -0.25 | 0.801 |
| 3 |  | 4 | uncorrected | common | recall | TSSeq | SVC | -0.01 | -0.10 | 0.921 |
| 3 |  | 4 | uncorrected | common | recall | HydraMultiROCKET | SVC | 0.00 | -0.05 | 0.959 |
| 3 |  | 4 | uncorrected | common | recall | CatEmbed | XGB | -0.01 | -0.14 | 0.891 |
| 3 |  | 4 | uncorrected | common | recall | GANDALF | XGB | 0.00 | -0.05 | 0.963 |
| 3 |  | 4 | uncorrected | common | recall | TabPFN | XGB | 0.00 | 0.03 | 0.975 |
| 3 |  | 4 | uncorrected | common | recall | LSTM | XGB | -0.02 | -0.22 | 0.829 |
| 3 |  | 4 | uncorrected | common | recall | TSSeq | XGB | 0.00 | -0.06 | 0.949 |
| 3 |  | 4 | uncorrected | common | recall | HydraMultiROCKET | XGB | 0.00 | -0.02 | 0.988 |
| 3 |  | 4 | uncorrected | common | recall | GANDALF | CatEmbed | 0.01 | 0.09 | 0.929 |
| 3 |  | 4 | uncorrected | common | recall | TabPFN | CatEmbed | 0.01 | 0.17 | 0.866 |
| 3 |  | 4 | uncorrected | common | recall | LSTM | CatEmbed | -0.01 | -0.08 | 0.937 |
| 3 |  | 4 | uncorrected | common | recall | TSSeq | CatEmbed | 0.01 | 0.07 | 0.942 |
| 3 |  | 4 | uncorrected | common | recall | HydraMultiROCKET | CatEmbed | 0.01 | 0.12 | 0.904 |
| 3 |  | 4 | uncorrected | common | recall | TabPFN | GANDALF | 0.01 | 0.08 | 0.937 |
| 3 |  | 4 | uncorrected | common | recall | LSTM | GANDALF | -0.01 | -0.17 | 0.866 |
| 3 |  | 4 | uncorrected | common | recall | TSSeq | GANDALF | 0.00 | -0.02 | 0.986 |
| 3 |  | 4 | uncorrected | common | recall | HydraMultiROCKET | GANDALF | 0.00 | 0.03 | 0.975 |
| 3 |  | 4 | uncorrected | common | recall | LSTM | TabPFN | -0.02 | -0.25 | 0.804 |
| 3 |  | 4 | uncorrected | common | recall | TSSeq | TabPFN | -0.01 | -0.10 | 0.924 |
| 3 |  | 4 | uncorrected | common | recall | HydraMultiROCKET | TabPFN | 0.00 | -0.05 | 0.962 |
| 3 |  | 4 | uncorrected | common | recall | TSSeq | LSTM | 0.01 | 0.15 | 0.879 |
| 3 |  | 4 | uncorrected | common | recall | HydraMultiROCKET | LSTM | 0.02 | 0.20 | 0.841 |
| 3 |  | 4 | uncorrected | common | recall | HydraMultiROCKET | TSSeq | 0.00 | 0.05 | 0.961 |
| 3 |  | 4 | uncorrected | uncommon | recall | SVC | RF | -0.38 | -6.07 | < 0.001 |
| 3 |  | 4 | uncorrected | uncommon | recall | XGB | RF | 0.02 | 0.23 | 0.816 |
| 3 |  | 4 | uncorrected | uncommon | recall | CatEmbed | RF | 0.00 | 0.01 | 0.992 |
| 3 |  | 4 | uncorrected | uncommon | recall | GANDALF | RF | -0.05 | -0.59 | 0.556 |
| 3 |  | 4 | uncorrected | uncommon | recall | TabPFN | RF | 0.02 | 0.30 | 0.768 |
| 3 |  | 4 | uncorrected | uncommon | recall | LSTM | RF | -0.21 | -2.83 | 0.005 |
| 3 |  | 4 | uncorrected | uncommon | recall | TSSeq | RF | -0.01 | -0.12 | 0.908 |
| 3 |  | 4 | uncorrected | uncommon | recall | HydraMultiROCKET | RF | 0.04 | 0.58 | 0.565 |
| 3 |  | 4 | uncorrected | uncommon | recall | XGB | SVC | 0.40 | 6.37 | < 0.001 |
| 3 |  | 4 | uncorrected | uncommon | recall | CatEmbed | SVC | 0.38 | 6.08 | < 0.001 |
| 3 |  | 4 | uncorrected | uncommon | recall | GANDALF | SVC | 0.34 | 5.35 | < 0.001 |
| 3 |  | 4 | uncorrected | uncommon | recall | TabPFN | SVC | 0.41 | 6.46 | < 0.001 |
| 3 |  | 4 | uncorrected | uncommon | recall | LSTM | SVC | 0.17 | 3.00 | 0.003 |
| 3 |  | 4 | uncorrected | uncommon | recall | TSSeq | SVC | 0.37 | 5.93 | < 0.001 |
| 3 |  | 4 | uncorrected | uncommon | recall | HydraMultiROCKET | SVC | 0.43 | 6.84 | < 0.001 |
| 3 |  | 4 | uncorrected | uncommon | recall | CatEmbed | XGB | -0.02 | -0.22 | 0.823 |
| 3 |  | 4 | uncorrected | uncommon | recall | GANDALF | XGB | -0.06 | -0.82 | 0.411 |
| 3 |  | 4 | uncorrected | uncommon | recall | TabPFN | XGB | 0.00 | 0.06 | 0.950 |
| 3 |  | 4 | uncorrected | uncommon | recall | LSTM | XGB | -0.23 | -3.08 | 0.002 |
| 3 |  | 4 | uncorrected | uncommon | recall | TSSeq | XGB | -0.03 | -0.35 | 0.727 |
| 3 |  | 4 | uncorrected | uncommon | recall | HydraMultiROCKET | XGB | 0.03 | 0.34 | 0.732 |
| 3 |  | 4 | uncorrected | uncommon | recall | GANDALF | CatEmbed | -0.05 | -0.60 | 0.550 |
| 3 |  | 4 | uncorrected | uncommon | recall | TabPFN | CatEmbed | 0.02 | 0.29 | 0.775 |
| 3 |  | 4 | uncorrected | uncommon | recall | LSTM | CatEmbed | -0.21 | -2.84 | 0.004 |
| 3 |  | 4 | uncorrected | uncommon | recall | TSSeq | CatEmbed | -0.01 | -0.13 | 0.900 |
| 3 |  | 4 | uncorrected | uncommon | recall | HydraMultiROCKET | CatEmbed | 0.04 | 0.57 | 0.571 |
| 3 |  | 4 | uncorrected | uncommon | recall | TabPFN | GANDALF | 0.07 | 0.89 | 0.376 |
| 3 |  | 4 | uncorrected | uncommon | recall | LSTM | GANDALF | -0.16 | -2.21 | 0.027 |
| 3 |  | 4 | uncorrected | uncommon | recall | TSSeq | GANDALF | 0.04 | 0.47 | 0.636 |
| 3 |  | 4 | uncorrected | uncommon | recall | HydraMultiROCKET | GANDALF | 0.09 | 1.17 | 0.243 |
| 3 |  | 4 | uncorrected | uncommon | recall | LSTM | TabPFN | -0.23 | -3.15 | 0.002 |
| 3 |  | 4 | uncorrected | uncommon | recall | TSSeq | TabPFN | -0.03 | -0.41 | 0.681 |
| 3 |  | 4 | uncorrected | uncommon | recall | HydraMultiROCKET | TabPFN | 0.02 | 0.28 | 0.779 |
| 3 |  | 4 | uncorrected | uncommon | recall | TSSeq | LSTM | 0.20 | 2.71 | 0.007 |
| 3 |  | 4 | uncorrected | uncommon | recall | HydraMultiROCKET | LSTM | 0.26 | 3.46 | 0.001 |
| 3 |  | 4 | uncorrected | uncommon | recall | HydraMultiROCKET | TSSeq | 0.05 | 0.69 | 0.489 |
| 3 |  | 4 | uncorrected | rare | recall | SVC | RF | -0.39 | -6.66 | < 0.001 |
| 3 |  | 4 | uncorrected | rare | recall | XGB | RF | 0.03 | 0.38 | 0.707 |
| 3 |  | 4 | uncorrected | rare | recall | CatEmbed | RF | -0.06 | -0.84 | 0.404 |
| 3 |  | 4 | uncorrected | rare | recall | GANDALF | RF | -0.17 | -2.28 | 0.023 |
| 3 |  | 4 | uncorrected | rare | recall | TabPFN | RF | 0.10 | 1.28 | 0.201 |
| 3 |  | 4 | uncorrected | rare | recall | LSTM | RF | -0.30 | -4.52 | < 0.001 |
| 3 |  | 4 | uncorrected | rare | recall | TSSeq | RF | -0.19 | -2.60 | 0.009 |
| 3 |  | 4 | uncorrected | rare | recall | HydraMultiROCKET | RF | 0.13 | 1.65 | 0.099 |
| 3 |  | 4 | uncorrected | rare | recall | XGB | SVC | 0.42 | 7.14 | < 0.001 |
| 3 |  | 4 | uncorrected | rare | recall | CatEmbed | SVC | 0.33 | 5.69 | < 0.001 |
| 3 |  | 4 | uncorrected | rare | recall | GANDALF | SVC | 0.22 | 4.25 | < 0.001 |
| 3 |  | 4 | uncorrected | rare | recall | TabPFN | SVC | 0.49 | 8.43 | < 0.001 |
| 3 |  | 4 | uncorrected | rare | recall | LSTM | SVC | 0.09 | 2.32 | 0.020 |
| 3 |  | 4 | uncorrected | rare | recall | TSSeq | SVC | 0.20 | 3.97 | < 0.001 |
| 3 |  | 4 | uncorrected | rare | recall | HydraMultiROCKET | SVC | 0.52 | 9.02 | < 0.001 |
| 3 |  | 4 | uncorrected | rare | recall | CatEmbed | XGB | -0.09 | -1.21 | 0.225 |
| 3 |  | 4 | uncorrected | rare | recall | GANDALF | XGB | -0.20 | -2.67 | 0.008 |
| 3 |  | 4 | uncorrected | rare | recall | TabPFN | XGB | 0.07 | 0.90 | 0.369 |
| 3 |  | 4 | uncorrected | rare | recall | LSTM | XGB | -0.33 | -4.96 | < 0.001 |
| 3 |  | 4 | uncorrected | rare | recall | TSSeq | XGB | -0.22 | -3.00 | 0.003 |
| 3 |  | 4 | uncorrected | rare | recall | HydraMultiROCKET | XGB | 0.10 | 1.27 | 0.206 |
| 3 |  | 4 | uncorrected | rare | recall | GANDALF | CatEmbed | -0.10 | -1.42 | 0.155 |
| 3 |  | 4 | uncorrected | rare | recall | TabPFN | CatEmbed | 0.16 | 2.13 | 0.033 |
| 3 |  | 4 | uncorrected | rare | recall | LSTM | CatEmbed | -0.23 | -3.60 | 0.000 |
| 3 |  | 4 | uncorrected | rare | recall | TSSeq | CatEmbed | -0.12 | -1.73 | 0.083 |
| 3 |  | 4 | uncorrected | rare | recall | HydraMultiROCKET | CatEmbed | 0.19 | 2.51 | 0.012 |
| 3 |  | 4 | uncorrected | rare | recall | TabPFN | GANDALF | 0.27 | 3.65 | 0.000 |
| 3 |  | 4 | uncorrected | rare | recall | LSTM | GANDALF | -0.13 | -2.15 | 0.032 |
| 3 |  | 4 | uncorrected | rare | recall | TSSeq | GANDALF | -0.02 | -0.31 | 0.759 |
| 3 |  | 4 | uncorrected | rare | recall | HydraMultiROCKET | GANDALF | 0.30 | 4.06 | < 0.001 |
| 3 |  | 4 | uncorrected | rare | recall | LSTM | TabPFN | -0.40 | -6.07 | < 0.001 |
| 3 |  | 4 | uncorrected | rare | recall | TSSeq | TabPFN | -0.29 | -3.99 | < 0.001 |
| 3 |  | 4 | uncorrected | rare | recall | HydraMultiROCKET | TabPFN | 0.03 | 0.36 | 0.717 |
| 3 |  | 4 | uncorrected | rare | recall | TSSeq | LSTM | 0.11 | 1.84 | 0.065 |
| 3 |  | 4 | uncorrected | rare | recall | HydraMultiROCKET | LSTM | 0.42 | 6.56 | < 0.001 |
| 3 |  | 4 | uncorrected | rare | recall | HydraMultiROCKET | TSSeq | 0.32 | 4.41 | < 0.001 |

##### Table S9:

The marginal means and 95% confidence intervals (CI_low_ – CI_high_) of the proportion of each behavioural class from the focal sampling and the predictions from the two different models (HR_B1_Unc: HydraMultiROCKET trained on burst 1 uncorrected dataset; Tab_B4_Bas: TabPFN trained on burst 4 dataset with basal rotation correction) in experiment 4.

| **Behaviour** | **Method** | **Value** | **CI_low_** | **CI_high_** |
| --- | --- | --- | --- | --- |
| Eating | Focal | 0.41 | 0.39 | 0.43 |
| Eating | HR_B1_Unc | 0.37 | 0.35 | 0.39 |
| Eating | Tab_B4_Bas | 0.35 | 0.33 | 0.37 |
| Resting | Focal | 0.26 | 0.24 | 0.28 |
| Resting | HR_B1_Unc | 0.33 | 0.31 | 0.35 |
| Resting | Tab_B4_Bas | 0.39 | 0.37 | 0.42 |
| Walking | Focal | 0.06 | 0.06 | 0.07 |
| Walking | HR_B1_Unc | 0.15 | 0.14 | 0.17 |
| Walking | Tab_B4_Bas | 0.14 | 0.12 | 0.15 |
| Grooming actor | Focal | 0.02 | 0.02 | 0.02 |
| Grooming actor | HR_B1_Unc | 0.06 | 0.05 | 0.06 |
| Grooming actor | Tab_B4_Bas | 0.05 | 0.04 | 0.06 |
| Grooming receiver | Focal | 0.03 | 0.02 | 0.03 |
| Grooming receiver | HR_B1_Unc | 0.10 | 0.09 | 0.11 |
| Grooming receiver | Tab_B4_Bas | 0.07 | 0.06 | 0.09 |
| Sleeping | Focal | 0.01 | 0.01 | 0.02 |
| Sleeping | HR_B1_Unc | 0.04 | 0.03 | 0.05 |
| Sleeping | Tab_B4_Bas | 0.04 | 0.04 | 0.05 |
| Self-scratching | Focal | 0.04 | 0.04 | 0.05 |
| Self-scratching | HR_B1_Unc | 0.01 | 0.01 | 0.01 |
| Self-scratching | Tab_B4_Bas | 0.02 | 0.02 | 0.03 |
| Running | Focal | 0.02 | 0.01 | 0.02 |
| Running | HR_B1_Unc | 0.02 | 0.02 | 0.03 |
| Running | Tab_B4_Bas | 0.02 | 0.02 | 0.02 |

##### Table S10:

The difference, z value and p value of post-hoc comparisons between various levels of the main predictors from the statistical comparisons for experiment 4.

| **Behaviour** | **Level1** | **Level2** | **Difference** | **z** | **p** |
| --- | --- | --- | --- | --- | --- |
| Eating | HR_B1_Unc | Focal | -0.04 | -2.43 | 0.015 |
| Eating | Tab_B4_Bas | Focal | -0.06 | -3.85 | 0.000 |
| Eating | Tab_B4_Bas | HR_B1_Unc | -0.02 | -1.42 | 0.156 |
| Resting | HR_B1_Unc | Focal | 0.07 | 4.67 | < 0.001 |
| Resting | Tab_B4_Bas | Focal | 0.13 | 8.89 | < 0.001 |
| Resting | Tab_B4_Bas | HR_B1_Unc | 0.06 | 4.18 | < 0.001 |
| Walking | HR_B1_Unc | Focal | 0.09 | 9.33 | < 0.001 |
| Walking | Tab_B4_Bas | Focal | 0.07 | 8.22 | < 0.001 |
| Walking | Tab_B4_Bas | HR_B1_Unc | -0.01 | -1.16 | 0.247 |
| Grooming actor | HR_B1_Unc | Focal | 0.03 | 7.04 | < 0.001 |
| Grooming actor | Tab_B4_Bas | Focal | 0.03 | 6.11 | < 0.001 |
| Grooming actor | Tab_B4_Bas | HR_B1_Unc | -0.01 | -1.09 | 0.276 |
| Grooming receiver | HR_B1_Unc | Focal | 0.07 | 10.79 | < 0.001 |
| Grooming receiver | Tab_B4_Bas | Focal | 0.05 | 8.00 | < 0.001 |
| Grooming receiver | Tab_B4_Bas | HR_B1_Unc | -0.03 | -3.22 | 0.001 |
| Sleeping | HR_B1_Unc | Focal | 0.03 | 7.19 | < 0.001 |
| Sleeping | Tab_B4_Bas | Focal | 0.03 | 7.87 | < 0.001 |
| Sleeping | Tab_B4_Bas | HR_B1_Unc | 0.00 | 0.92 | 0.360 |
| Self-scratching | HR_B1_Unc | Focal | -0.03 | -7.92 | < 0.001 |
| Self-scratching | Tab_B4_Bas | Focal | -0.02 | -4.35 | < 0.001 |
| Self-scratching | Tab_B4_Bas | HR_B1_Unc | 0.01 | 4.50 | < 0.001 |
| Running | HR_B1_Unc | Focal | 0.00 | 1.56 | 0.119 |
| Running | Tab_B4_Bas | Focal | 0.00 | 1.13 | 0.258 |
| Running | Tab_B4_Bas | HR_B1_Unc | 0.00 | -0.44 | 0.663 |
